# Knock-in *Kcnh2* Rabbit Model of Long QT Syndrome Type-2, Epilepsy, and Sudden Death

**DOI:** 10.1101/2024.12.11.627988

**Authors:** Veronica Singh, Kyle T. Wagner, Laura G. Williams, Justin M. Ryan, Katherine R. Keller, Jonathan D. Mohnkern, Robert S. Gardner, Louis T. Dang, Julie M. Ziobro, Richard J. H. Wojcikiewicz, Nathan R. Tucker, David S. Auerbach

## Abstract

**Background:** Long QT Syndrome Type-2 (LQT2) is due to loss-of-function *KCNH2* variants. *KCNH2* encodes K_v_11.1 that forms a delayed-rectifier potassium channel in the brain and heart. LQT2 is associated with arrhythmias, seizures, sudden cardiac death, and sudden unexpected death in epilepsy (SUDEP). The goal of the study is to develop a translational model that reproduces the neuro-cardiac electrical abnormalities and sudden death seen in people with LQT2.

**Methods:** We generated the first knock-in rabbit model of LQT2 (*Kcnh2*^(+/7bp-del)^), due to a 7 base-pair (7bp) deletion in the pore domain of the endogenous rabbit *Kcnh2* gene.

**Results:** Mutant *Kcnh2* is expressed in the heart and brain and constitutes 11% of total *Kcnh2* in *Kcnh2*^(+/7bp-del)^ rabbits. Total *Kcnh2*, WT *Kcnh2*, and WT K_v_11.1 expression is lower in *Kcnh2*^(+/7bp-del)^ vs. WT rabbits. *Kcnh2*^(+/7bp-del)^ rabbits exhibit prolonged cardiac ventricular repolarization (QT_c_, JT_ec_, JT_pc_). There is an increased prevalence of spontaneous epileptiform activity and clinical seizures in *Kcnh2*^(+/7bp-del)^ (7 of 37 rabbits) vs. WT rabbits (1:68 rabbits, *p*<0.003). 18.9% of *Kcnh2*^(+/7bp-del)^ vs. 1.5% of WT rabbits died suddenly and spontaneously (*p*<0.003). We recorded 2 spontaneous lethal events in *Kcnh2*^(+/7bp-del)^ rabbits: (1) sudden cardiac death and (2) seizure-mediated sudden death due to generalized tonic-clonic seizures, post-ictal generalized EEG suppression, bradycardia, ECG-T-wave inversion, focal cardiac activity, and asystole/death.

**Conclusions:** We developed the first genetic rabbit model of LQT2 that reproduces the cardiac and epileptic phenotypes seen in people with LQT2. *Kcnh2*^(+/7bp-del)^ rabbits provide a valuable tool for future mechanistic studies, development of neurotherapeutics, and cardiac-safety testing.

## Introduction

Long QT Syndrome (LQTS) is an ion channelopathy associated with a high risk of cardiac arrhythmias (e.g., *torsade de pointes*) and sudden cardiac death (SCD)(1). It affects 1:2000 people and is characterized by prolongation of the cardiac electrical activation-recovery interval (QT_c_ on the electrocardiogram, ECG)(2, 3). Long QT Syndrome Type-2 (LQT2) is due to loss-of-function (LOF) variants in the *KCNH2* gene(4, 5). *KCNH2* encodes the K_v_11.1 protein, which forms the α- subunit of the channel that passes the rapid delayed rectifier potassium current (I_Kr_)(6). I_Kr_ is responsible for cardiomyocyte repolarization(7) and suppression of repetitive action potential (AP) firing in neurons(8). *KCNH2* LOF variants cause a reduction in I_Kr_, leading to cardiomyocyte AP prolongation and hyperexcitability, which ultimately provides a substrate for arrhythmias and SCD. Interestingly, there is a 3.7-fold higher prevalence of seizures in people genotype-positive for LQT2, compared to genotype-negative family members(9). The prevalence of electroencephalogram (EEG) diagnosed epilepsy is higher in people with LQT2, compared to people with LQTS Type-1 (LQT1) and healthy controls without epilepsy(10, 11). Post-mortem genetic analysis indicates a higher prevalence of loss-of-function (3-fold) and rare (11-fold) *KCNH2* variants in Sudden Unexpected Death in Epilepsy (SUDEP) cases vs. living epilepsy controls(12). Despite overwhelming evidence of a dual pathological role of *KCNH2* variants in the heart and brain, there is no translational model that fully reproduces the neuro-cardiac abnormalities seen in people with LQT2. There is an unmet need for a clinically relevant model of LQT2 to investigate the mechanisms for the high risk of seizures and SUDEP in LQT2 and *KCNH2*-mediated epilepsy.

The objective of this study is to generate and characterize a genetic rabbit model of *Kcnh2*-mediated epilepsy, ECG abnormalities, and sudden death (SCD & SUDEP), which reproduces the neuro-cardiac electrical abnormalities seen in people with LQT2. We developed the first knock-in rabbit model of LQT2 that is due to a heterozygous frameshift deletion mutation in the pore domain of the endogenous rabbit *Kcnh2* gene. *KCNH2* pore variants confer the highest risk of arrhythmias and seizures in people with LQT2(1, 9). A novel clinically relevant animal model of LQT2 will facilitate future studies to investigate the underlying mechanisms of EEG abnormalities, epileptic seizures, and SUDEP, as well as drug development and testing.

## Methods

All experiments were performed in accordance with the Guide for the Care and Use of Laboratory Animals and approved by the Institutional Animal Care and Use Committee.

*Generation of Knock-In Rabbit Model of LQT2:* Using CRISPR-Cas9 technology, the Center for Advanced Models and Translational Sciences and Therapeutics (CAMTraST) at the University of Michigan generated a founder rabbit (*Kcnh2*^(+/7bp-del)^) with a 7bp frameshift deletion (1627bp-1633bp, NM_001082384.1) in one allele of the endogenous rabbit *Kcnh2* gene. *Kcnh2*^(+/7bp-del)^ rabbits are cross-bred and maintained on the New Zealand White background (Charles River, Wilmington, Massachusetts). Rabbits >1-month of age are housed in separate cages at 22°C, fed *ad libitum*, and on a 12- hour light/dark cycle (lights on 6AM–6PM).

*RNA Sequencing, Oxford Nanopore Technology (ONT) Sequencing, and Quantitative PCR (qPCR)*: RNAseq libraries were constructed from rabbit left ventricle and brain stem samples using Zymo-Seq RiboFree Total RNA library kit and sequenced on an Illumina NovaseqX 10B flow cell. ONT sequencing was performed by Plasmidsaurus using primers that surround the mutation. For qPCR, PrimeTime probes (Integrated DNA Technologies) are designed using sequences specific to WT and mutant *Kcnh2* transcripts. Forward and reverse primers are also generated for beta-actin (Actb), which serves as the housekeeping gene. See supplement for details (Supplement Table 1).

*K_v_11.1 Expression*: We verified the K_v_11.1 expression in HEK cells and rabbit tissue using commercially available antibodies (anti-MYC, anti-FLAG) and an antibody generated in-house (anti-K_v_11.1). See supplement for details.

*In-Vivo Video-ECG-EEG Signal:* Cardiac (ECG), neuronal (EEG), audio, and video recordings are acquired from conscious restrained rabbits. Full details about the type of electrodes, EEG/ECG locations, and acquisition system are described in the supplement and in Bosinski *et al*. 2021(13). The electrode placements faciliate unipolar, bipolar, and referential EEG montages, as well as bipolar, augmented, and referential ECG configurations (Supplementary Fig. 1).

*ECG and EEG Analysis:* All ECG data is analyzed using LabChart8 and average representative genotype-specific ECG traces are generated using MATLAB 2023b. From each rabbit, a 5-minute baseline period of ECG signal during a stable heart rate is analyzed. All video/EEG recordings are manually reviewed by an investigator blinded to the genotype and sex, and all epileptiform activity and seizures are confirmed by board-certified pediatric epileptologists (LTD & JMZ). A 60 Hz notch filter and 1-70Hz bandpass filter is applied to all EEG signal.

## Results

*CRISPR-Generated Knock-In LQT2 Rabbit Model*: *Kcnh2*^(+/7bp-del)^ rabbits were generated with a 7bp deletion in the S5 pore domain of the rabbit endogenous *Kcnh2* gene (Fig. 1A), which creates 17 predicted premature stop codons. Mutant gDNA is present in tissue collected from the offspring of *Kcnh2*^(+/7bp-del)^, but not WT rabbits (Supplementary Fig. 2A). The mutation is germline and the *Kcnh2*^(+/7bp-del)^ offspring are viable and fertile. Despite multiple rounds of breeding 2 *Kcnh2*^(+/7bp-del)^ rabbits together, there were only *Kcnh2*^(+/7bp-del)^ and WT offspring; there were no *Kcnh2*^(7bp-del/7bp-del)^ rabbits.

**Fig. 1:**
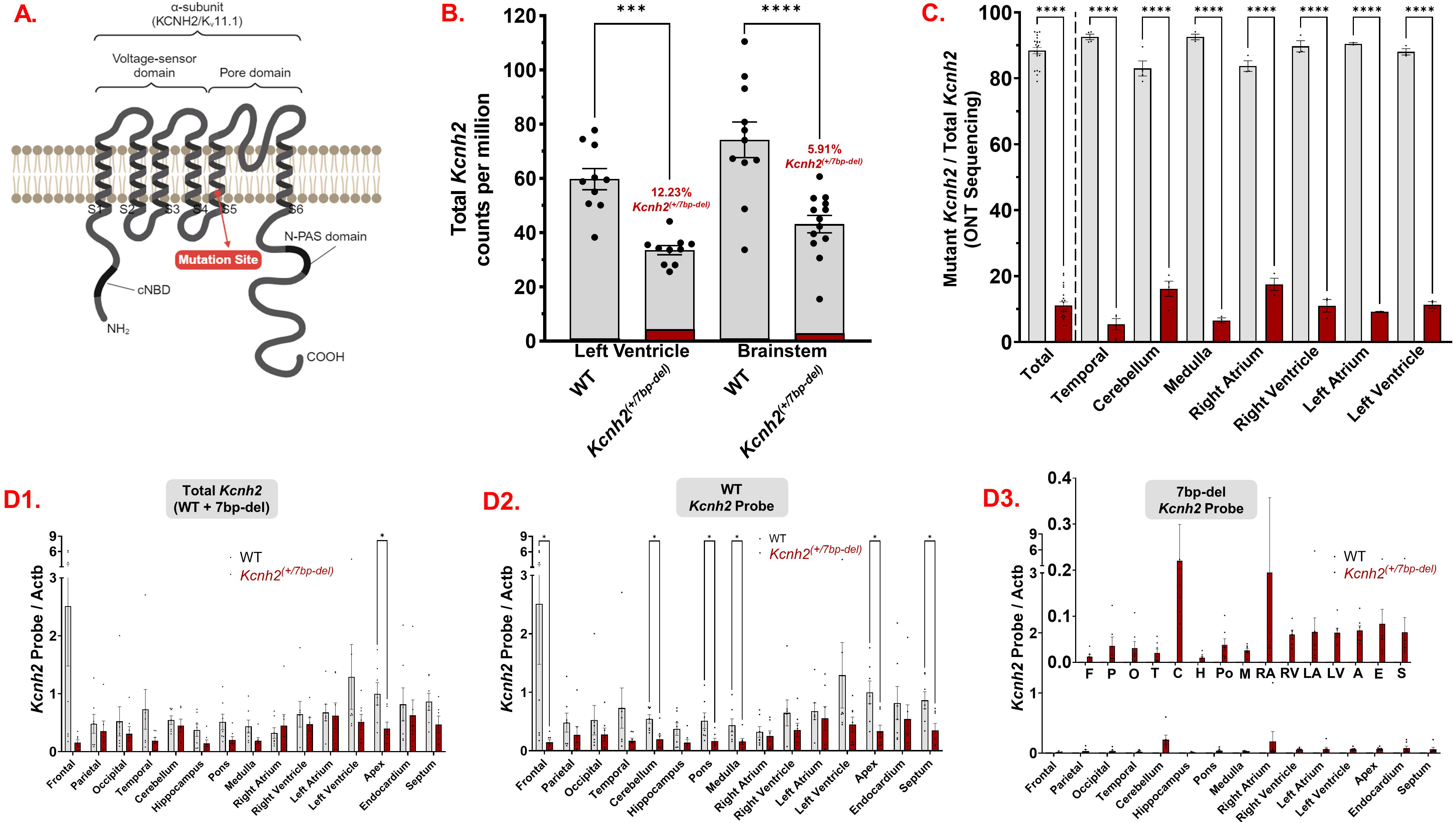
CRISPR Cas-9-mediated *Kcnh2* mutation leads to altered Kcnh2 expression patterns. **A.** Topology of K_v_11.1 protein denoting the site of 7bp-deletion mutation in S5 of the pore domain. **B.** RNA sequencing indicates altered total *Kcnh2* expression in WT and *Kcnh2*^(+/7bp-del)^ rabbit heart and brain tissue. WT left ventricle N=10 rabbits, WT brainstem N=11 rabbits, *Kcnh2*^(+/7bp-del)^ left ventricle N=10 rabbits, *Kcnh2*^(+/7bp-del)^ brainstem N=13 rabbits. **C.** Oxford Nanopore Technology (ONT) sequencing of WT and mutant *Kcnh2* amplicons generated using RNA extracted from mutant rabbits (N=3). **D.** qPCR resuls for region-specific expression of **1.** Total (WT+mutant), **2.** WT, and **3.** 7bp-del mutant *Kcnh2* transcripts in WT (N=7) vs. mutant (N=7) rabbit tissue. All qPCR data is normalized to plate normalization factor (*described in results*). *, p<0.05; **, *p*<0.01; ****, *p*<0.0001. Actb: beta-actin.

*WT/Mutant Kcnh2 Expression Patterns*: RNA sequencing results indicate a 43.9% reduction of total *Kcnh2* in the heart (left ventricle) and 41.9% reduction in the brain (brainstem) in *Kcnh2*^(+/7bp-del)^ vs. WT tissue (Fig. 1B). In *Kcnh2*^(+/7bp-del)^ rabbits, mutant transcript comprises 12.2% and 5.9% of total *Kcnh2* transcript in the heart and brain, respectively. Similarly, ONT sequencing results demonstrate that mutant transcript is 11% of the total *Kcnh2* (Fig. 1C, range 5.4-16.1% in specific brain & heart regions).

We profiled tissue-specific total, WT, and mutant *Kcnh2* expression using qPCR. The primers amplify the mutant region in both WT and mutant tissue, and the primers/probes are specific to WT and mutant cDNA (Supplementary Figs. 2 & 3). Results from qPCR indicate that total and WT *Kcnh2* is lower in each region of the brain and heart of *Kcnh2*^(+/7bp-del)^ vs.

WT rabbits (Fig 1D1-2). In *Kcnh2*^(+/7bp-del)^ rabbits, total *Kcnh2* is significantly lower in the heart apex, and WT *Kcnh2* transcript is significantly lower in the frontal cortex, cerebellum, pons, and medulla of the brain, as well as the heart apex and septum. Mutant *Kcnh2* is detected in each of the brain and heart regions, with the highest expression in the cerebellum and right atrium (Fig. 1D3). In summary, results from RNA sequencing, ONT sequencing, and qPCR demonstrate that total and WT *Kcnh2* expression are each significantly lower in *Kcnh2*^(+/7bp-del)^, compared to WT tissue. Mutant *Kcnh2* comprises a small percentage of the total *Kcnh2* in *Kcnh2*^(+/7bp-del)^ tissue.

*K_v_11.1 Protein Expression:* We raised an antibody in guinea pigs against a peptide corresponding to the C-terminus of K_v_11.1 (Supplementary Table 1). The antibody is immunoreactive against full-length mammalian K_v_11.1^WT^. It recognizes N-terminally tagged exogenous rabbit MYC-K_v_11.1^WT^ at ∼145kDa, but not FLAG-Kv11.1^7bp-del^ in HEK cells (Supplementary Fig. 4A). Anti-MYC yields only a band at ∼145kDa in MYC-Kv11.1^WT^ . Consistent with the 7bp frameshift deletion creating predicted premature stop codons, a ∼72kDa band is detected in the FLAG-K_v_11.1^7bp-del^ samples using anti-FLAG, but not anti-K_v_11.1 (C-terminal epitope). This K_v_11.1 antibody also detects endogenous K_v_11.1^WT^ in mouse pituitary and human neuroblastoma cell lines (α-T3 & SH-SY5Y, Supplementary Fig 4B).

Membrane preparations from the heart and brain of WT and *Kcnh2*^(+/7bp-del)^ rabbits, indicate immunoreactive bands at 145- 150kDa (Fig 2A). K_v_11.1 expression is markedly lower in the left ventricle (Fig. 2A1) and temporal lobe (Fig. 2A2) of *Kcnh2*^(+/7bp-del^) vs. WT rabbit samples (n=3 rabbits/genotype). Interestingly, K_v_11.1 expression levels vary in specific regions of the WT brain (highest in cerebellum, brainstem and pons), but is consistently lower in *Kcnh2*^(+/7bp-del)^ tissue (Fig. 2B1). K_v_11.1 expression is consistenty lower in each of the regions of the heart of *Kcnh2*^(+/7bp-del)^ vs. WT rabbits (Fig. 2B2). Overall, full-length K_v_11.1 expression is lower throughout the brain and heart of *Kcnh2*^(+/7bp-del)^ vs. WT rabbits.

**Fig. 2:**
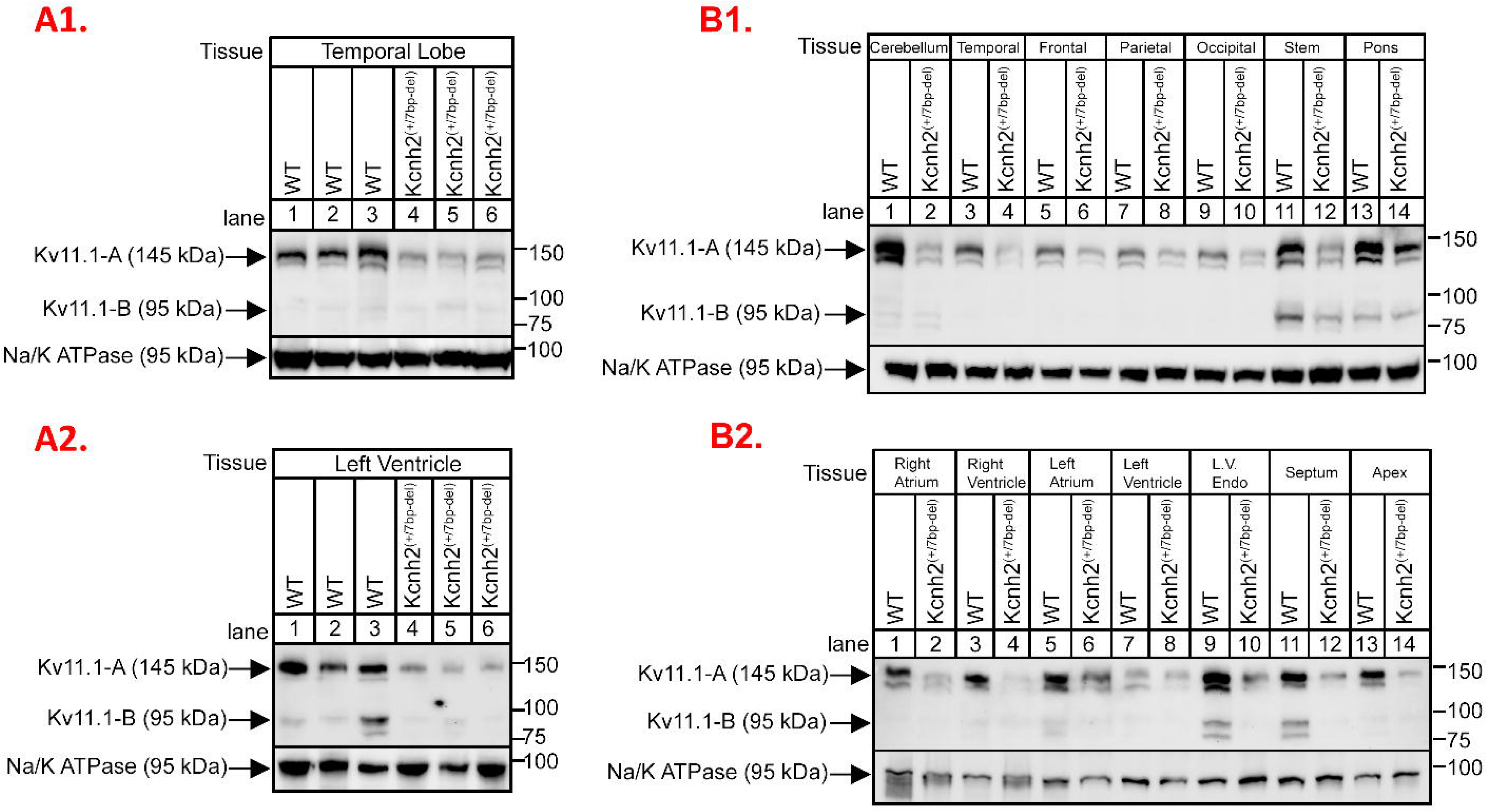
K_v_11.1 expression in rabbit tissue. **A.** Immunoblot showing immunoreactivity of C-terminus-specific K_v_11.1 antibody recognizing full-length K_v_11.1^WT^ in WT (N: 3) and *Kcnh2*^(+/7bp-del)^ (N: 3) **1.** heart (left ventricle) and **2.** brain (temporal lobe) tissue. **B.** Immunoblot showing region-specific expression of full-length K_v_11.1^WT^ in WT and *Kcnh2*^(+/7bp-del)^ **1.** heart and **2.** brain tissue.

*In Vivo Characteriztion of Cardiac Electrical Function:* Baseline conscious restrained video/EEG/ECG recordings were performed in WT and *Kcnh2*^(+/7bp-del)^ rabbits. Fig. 3A and B are 40-beat average ECG waveforms from WT and *Kcnh2*^(+/7bp-del)^ rabbits. In contrast to rodents(14), it resembles the human ECG waveform, as seen by positive and broad T-waves. The widely-split and notched (bifid), and prolonged T-wave morphology in the *Kcnh2*^(+/7bp-del)^ ECG trace reproduces the T-wave morphology seen in people with LQT2(15–17).

**Fig. 3:**
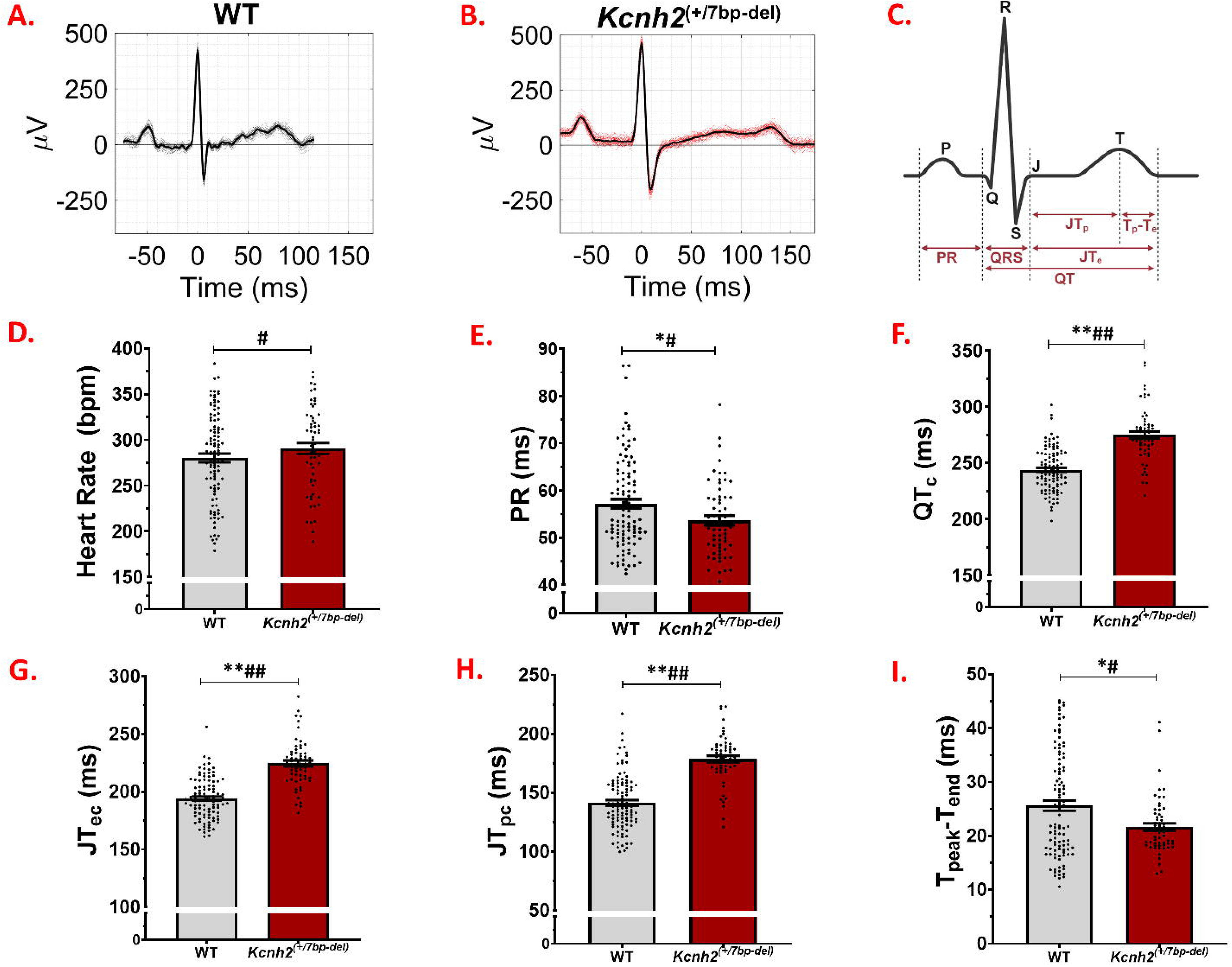
*In-vivo* cardiac phenotype in LQT2 knock-in rabbits. Representative ECG traces of **A.** WT (black) and **B.** *Kcnh2*^(+/7bp-del)^ (red) rabbits, showing the P, QRS, and T waves. **C.** Schematic representing ECG intervals that are quantified. Depolarization and repolarization metrics in WT (grey, N=68 rabbits, n=89 recordings) and *Kcnh2*^(+/7bp-del)^ (red, N=37 rabbits, n=52 recordings) rabbits: **D.** Heart rate, **E.** PR, **F.** QT_c_, **G.** JT_ec_, **H.** T_pc_, and **I.** T_peak_-T_end_. *, *p*<0.05; ** *p*<0.0001 Wilcoxon rank-sum test. ^#^, *p*<0.05; ^##^, *p*<0.0001 logistic regression models adjusted for age, sex, and heart rate.

We assessed heart rate, conduction, and repolarization metrics in WT vs. *Kcnh2*^(+/7bp-del)^ rabbits (Fig. 3C-I, Supplementary Table 2). P and QRS durations are similar in WT vs. *Kcnh2*^(+/7bp-del)^ rabbits. Atrio-ventricular conduction time (PR interval) is shorter in *Kcnh2*^(+/7bp-del)^ vs. WT Rabbits (*p*=0.018, Fig. 3E). Heart rate corrected repolarization measures, which include QT_c_ (*p*<0.001, Fig 3F), JT_ec_ (*p*<0.001, Fig 3G), and JT_pc_ (*p*<0.001, Fig 3H), are longer in *Kcnh2*^(+/7bp-del)^ (N=37 rabbits, n=52 recordings),vs. WT rabbits (N=68 rabbits, n=89 recordings). The T_peak_T_end_ is shorter in *Kcnh2*^(+/7bp-del)^ vs. WT rabbits (p<0.05, Fig 3I), which indicates reduced spatial and transmural dispersion of repolarization(18, 19). Similar results were also seen when stratifying for age (Supplementary Figure 5). Logistic regression models adjusting for age, sex, and heart rate further confirm that PR, QT_c_, JT_ec_, and JT_pc_ are prolonged, and T_peak_T_end_ is shorter in *Kcnh2*^(+/7bp-del)^ vs. WT rabbits. In summary, *Kcnh2*^(+/7bp-del)^ rabbits reproduce the human LQT2 ECG pathology of singificant prolongation of cardiac ventricular repolarization metrics(QT_c_, JT_pc_, & JT_ec_).

*In Vivo Characterization of EEG Recordings*: EEG recordings were manually reviewed to identify epileptiform activity and seizures (N=105 rabbits, n=225 30-60 minutes recordings). We followed the criteria for inter-ictal epileptiform discharges set by the American Academy of Neurology(20). Supplementary Fig. 1 shows a normal EEG/ECG recording from a 1.5- month-old male WT rabbit. We identified several instances of spontaneous epileptiform activity in *Kcnh2*^(+/7bp-del)^ rabbits. Interestingly, all of the cases of epileptiform activtiy and seizures are noted in juvenile rabbits (2-8 weeks of age). Fig. 4 shows an EEG recording from a 2-week-old male *Kcnh2*^(+/7bp-del)^ rabbit; it illustrates spontaneous epileptiform activity and an electrographic seizure. The red arrowindicates the start of the seizure, followed by the temporal evolution of epileptiform discharges that originate in the right occipital region, increase in amplitude, and are then seen in the left occipital leads. In a 7-week-old female *Kcnh2*^(+/7bp-del)^ rabbit, spontaneous epileptiform discharges and clonic head movement is noted (Fig. 5A). In a 4-week-old female *Kcnh2*^(+/7bp-del)^ rabbit, spike and slow wave epileptiform activity is noted in between myoclonic head jerks (Fig. 5B). In summary, spontaneous epileptiform activity and seizures are identified in 7 of 37 *Kcnh2*^(+/7bp-del)^ vs. 1 of 68 WT rabbits (Fig. 5C; p<0.01).

**Fig. 4:**
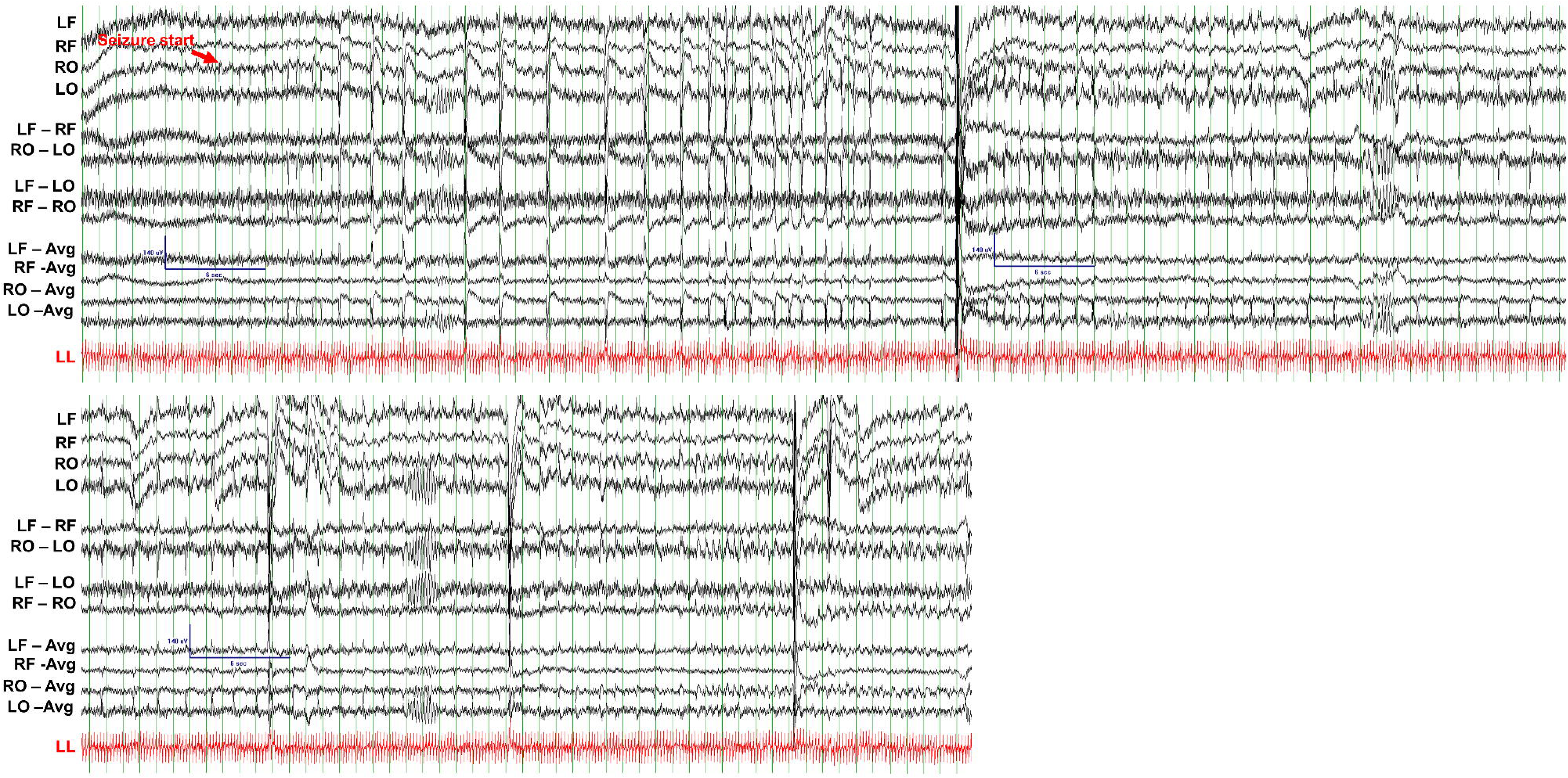
Representative EEG (black) and ECG (red) traces during an electrographic seizure in a 2-week-old male *Kcnh2*^(+/7bp-del)^ rabbit; epileptiform discharges in right occipital region increasing in amplitude over time. Red arrow indicates seizure start. LF: left frontal, RF: right frontal, RO: right occipital, LO: left occipital, LL: left leg, RA: right arm, LA: left arm. Scale bar shown for EEG signal: 140µV amplitude, 6 seconds.

**Fig. 5:**
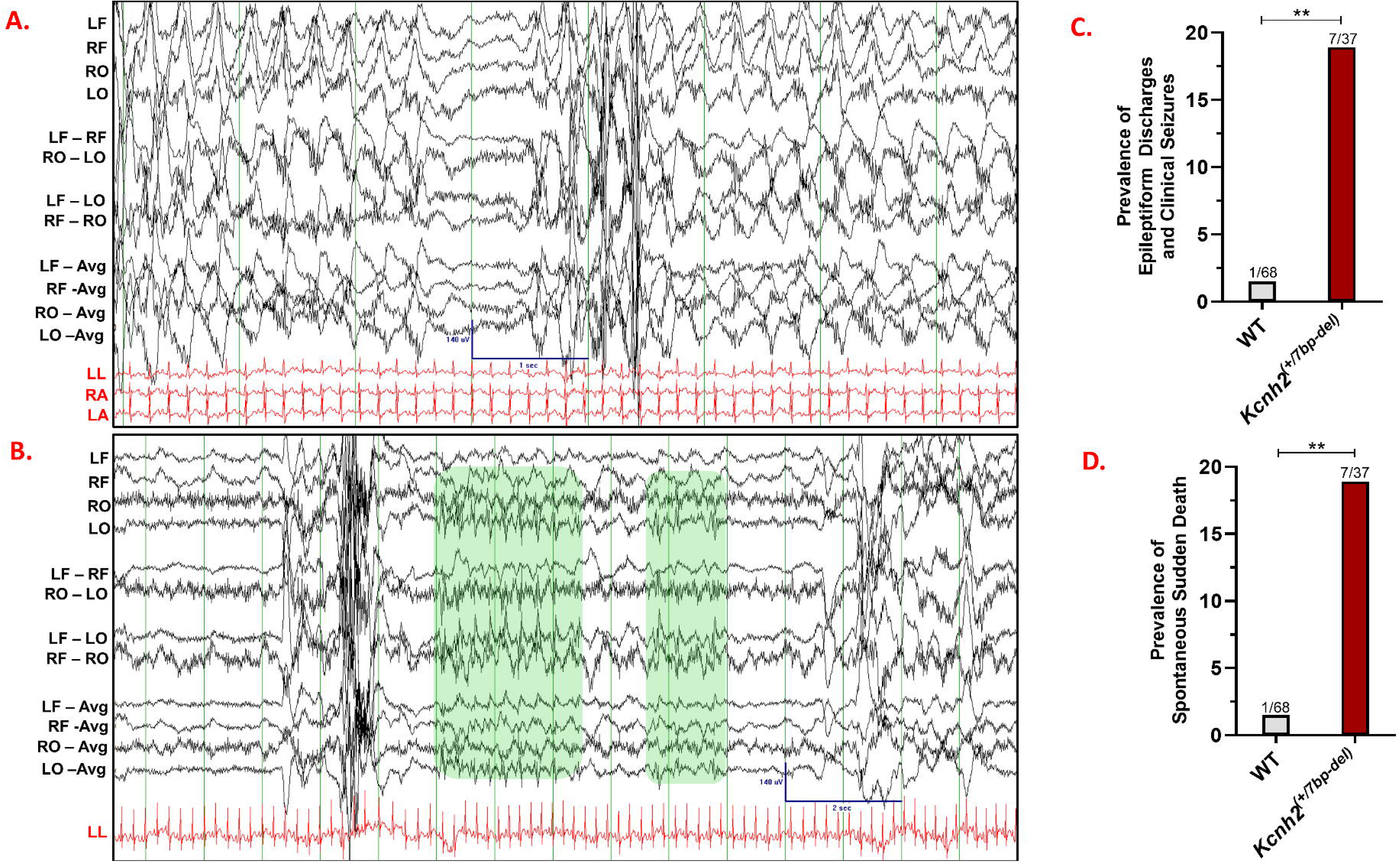
*In-vivo* neuronal phenotype in WT and *Kcnh2*^(+/7bp-del)^ rabbits. **A.** Spontaneous epileptiform discharges and a clonic seizure noted in a 7-week-old female *Kcnh2*^(+/7bp-del)^ rabbit. EEG scale bar indicates 140µV and 1 second. **B.** EEG-ECG traces between myoclonic jerks showing epileptiform discharges with a spike and slow wave morphology. EEG scale bar indicates 140µV and 2 seconds. **C.** Prevalence of epileptiform discharges and clinical seizures (WT N=68, n=140; *Kcnh2*^(+/7bp-del)^ N=37, n=85). **D.** Prevalence of spontaneous sudden death in WT (N=68) and *Kcnh2*^(+/7bp-del)^ (N=37) rabbits. **, *p*<0.01. N=number of animals, n=number of recordings. LF: left frontal, RF: right frontal, RO: right occipital, LO: left occipital, LL: left leg, RA: right arm, LA: left arm.

*Increased Prevalence of Spontaneous Sudden Death in Kcnh2*^(+/7bp-del)^ *Rabbits:* There is a 13-fold higher prevalence of spontaneous sudden death in *Kcnh2*^(+/7bp-del)^ (7 of 37 rabbits), compared to WT rabbits (1 of 68 rabbits, Fig. 5D; p<0.01). During necropsy, all of the rabbits were in rigor, despite 5 of the deaths either being witnessed or the rabbit was last seen alive ≤2 hours prior. We did not identify an apparent cause of death during external examination in the cage or during the necropsy. We ruled out structural heart disease, trauma or internal bleeding, overt infection, tumor, and gastrointestinal blockage. There was no sudden large weight change noted prior to death. Death occurred between the ages of 0.4 to 30 months of age. Sudden death was witnessed/videoed for 3 *Kcnh2*^(+/7bp-del)^ (Sudden Death Cases 1-3) and 1 WT rabbit (Sudden Death Case 8), and there are video/EEG/ECG recordings for Sudden Death Cases 1 and 2 (Table 1).

**Table 1:**
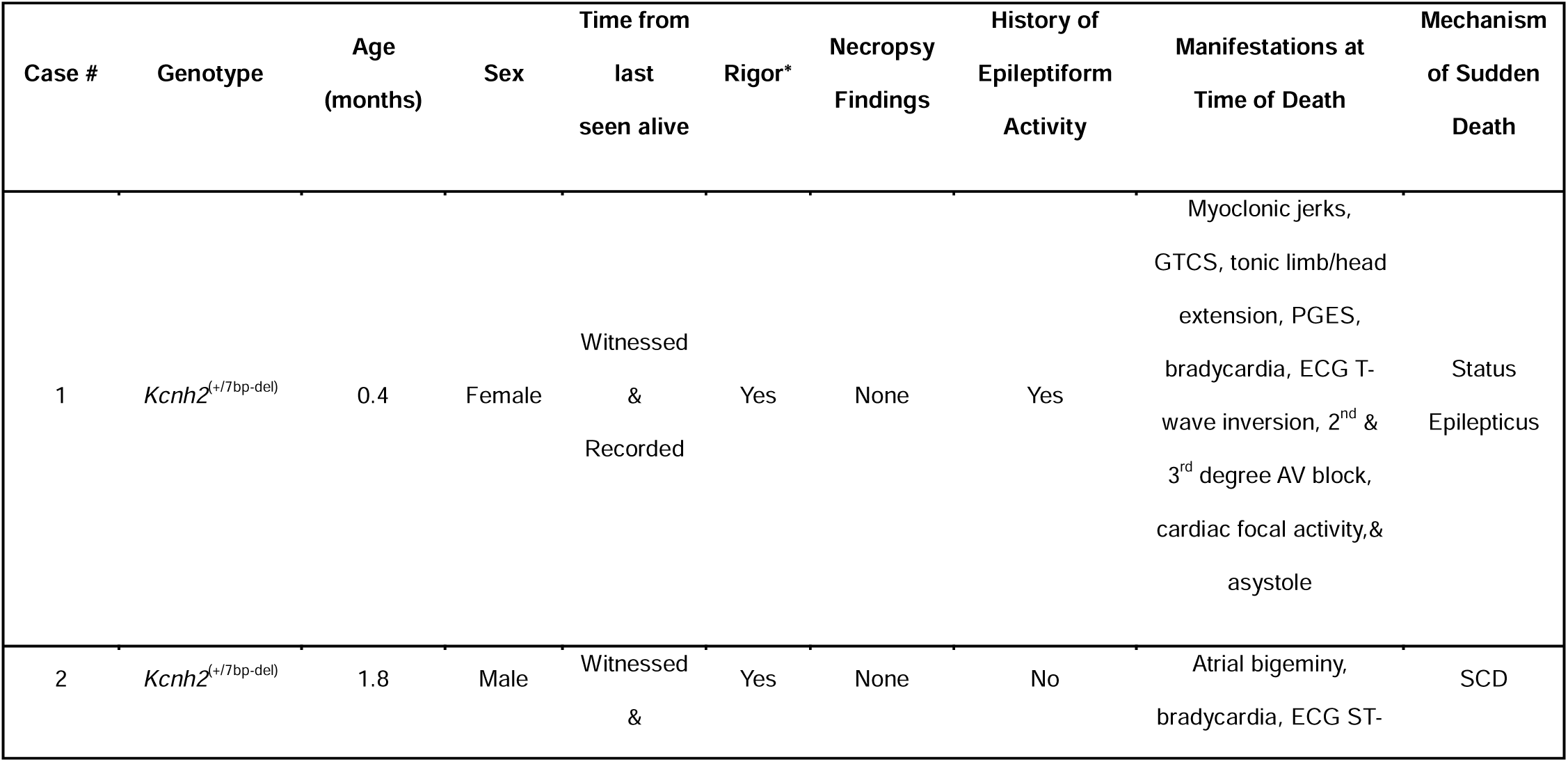
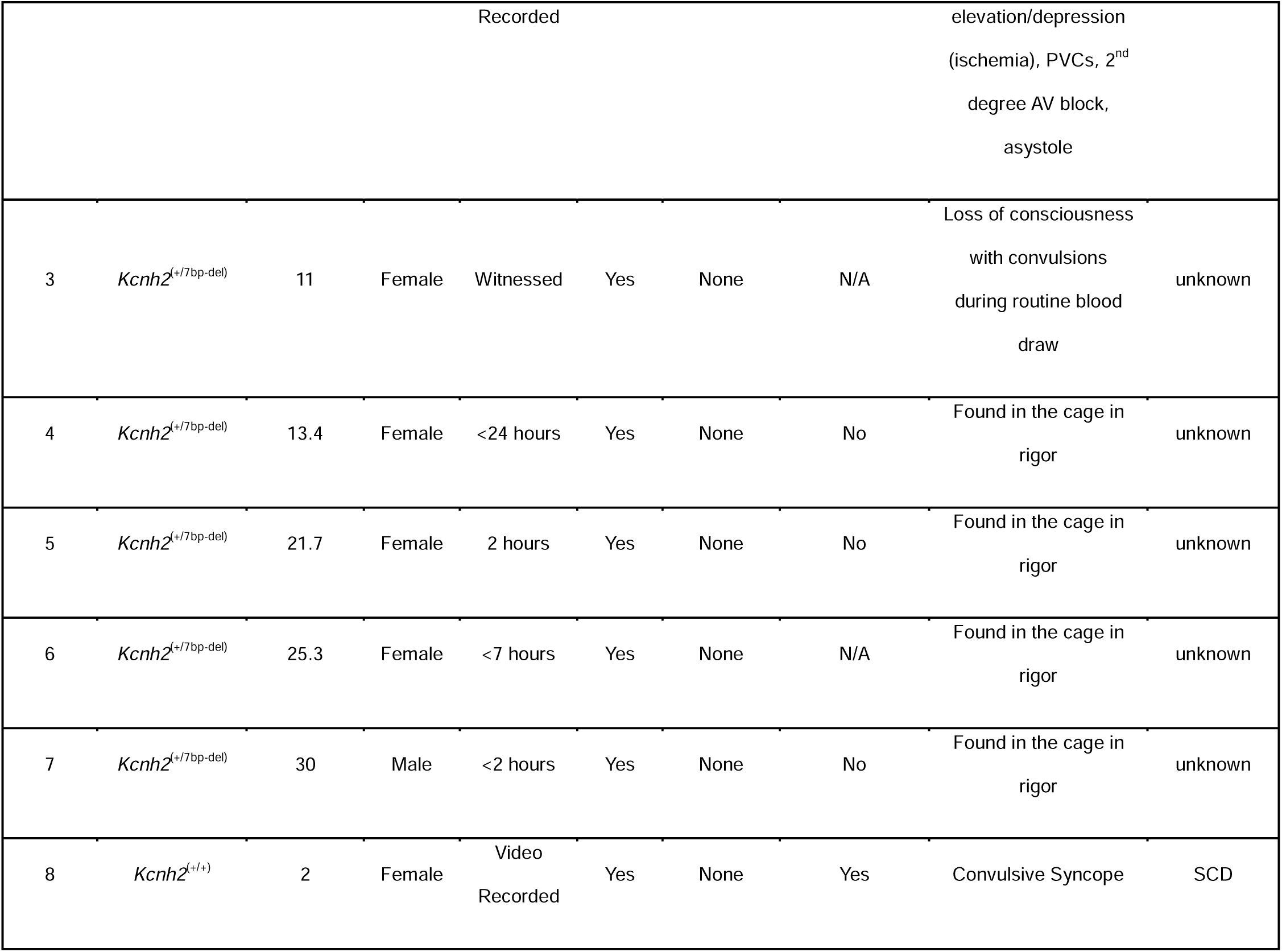
Detailed description of all cases of spontaneous sudden death. *\**: All animals that died suddenly and spontaneously were found in rigor, some within 2 hours of last being seen alive. Post-mortem necropsy did not reveal any anatomical or toxicological cause of death in any case of sudden death. SCD: sudden cardiac death.

Sudden Death Case 1 succomed to status-epilepticus-mediated sudden death. We examined the prevalence of epileptiform activity. Recordings acquired 24 and 48 hours prior to death indicate the presence of epileptiform activity. 2 of 3 mutant littermates also exhibited epileptiform activity, while no EEG abnormalities were noted in the 2 WT littermates. In this litter, the QT_c_ is longer in the 4 mutant vs. 2 WT rabbits (p<0.0001, Supplementary Table 3A). Interestingly, the inter- ictal QT_c_ for Sudden Death Case 1 is longer on the day of sudden death, compared to inter-ictal recordings 24 and 48 hours prior to sudden death (p<0.0001, Supplementary Table 3B). During a seizure, the QT_c_ prologation was so extreme that the P-wave is often hidden in the T wave of the previous beat (Fig. 6).

**Fig. 6:**
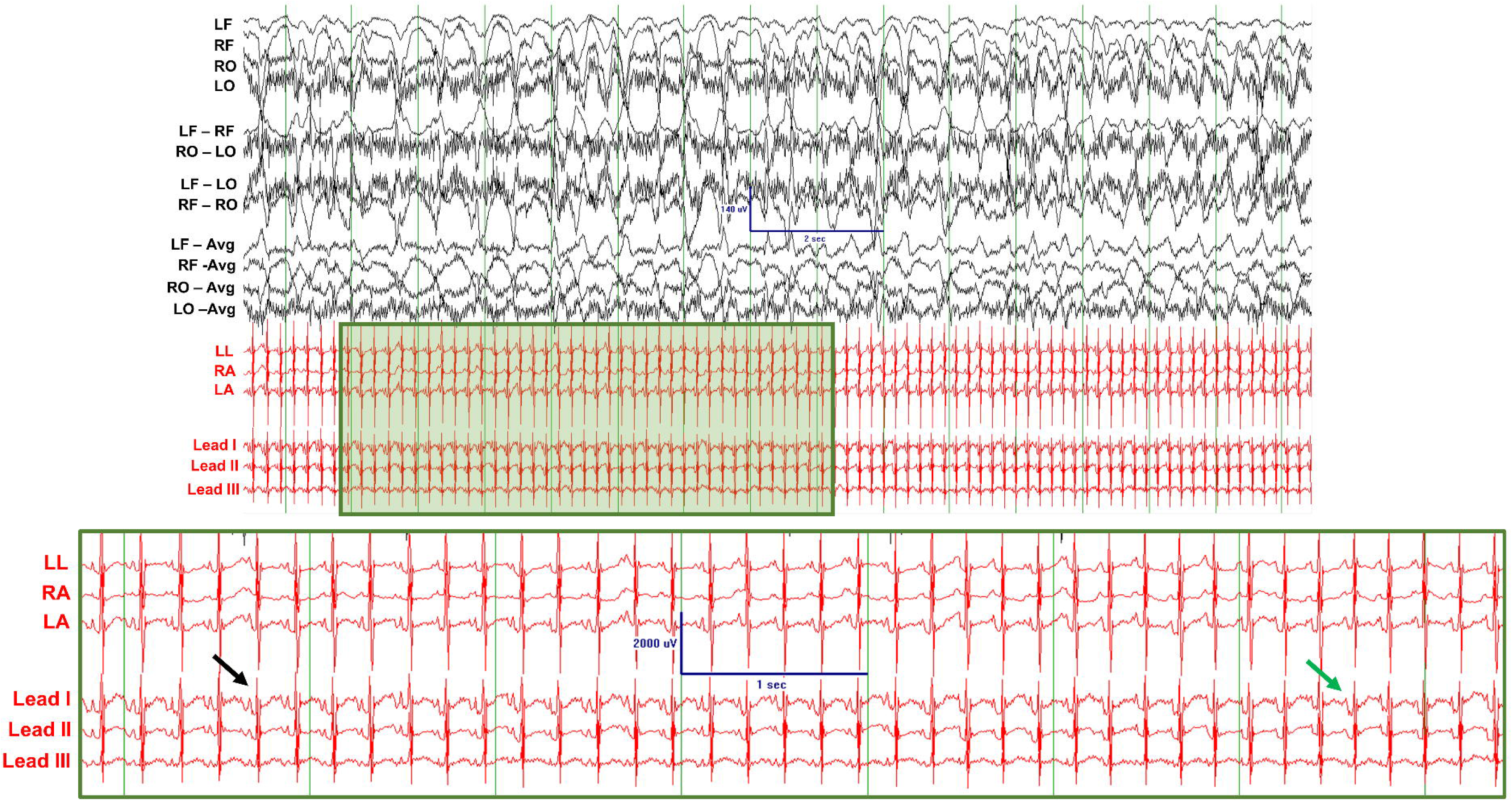
Sudden Death Case-1 exhibiting epileptiform discharges and extreme QT prolongation. Black arrow indicates a region where T and P waves of adjacent beats are temporally separated. Green arrow indicates a region where, due to extreme QT prolongation, the P wave is encapsulated by the T wave of a previous beat, causing an absence of the T-P interval. EEG scale bar indicates 140µV amplitude and 2 seconds; inset: ECG scale bar indicates 2000µV amplitude and 1 second.

On the day of sudden death, there are numerous instances of epileptiform activity, electrographic seizures, and epileptic seizures with profound clinical manifestations. The rabbit developed seizure clusters that included focal myoclonic seizures that transitioned to generalized tonic-clonic seizures (GTCS), which included periods of tonic limb extension and rhythmic movement of all extremities. Fig. 7A shows rhythmic ∼1.5Hz epileptiform discharges during a period that the rabbit presented with slight tremors. Amidst a background of myogenic artifact, there are EEG spikes that are higher in amplitude than the background, and the spikes have fast upstrokes followed by slower downstrokes. Following a particularly severe GTCS, there was a period of transient asystole lasting approximately 10 seconds (Fig. 7B1). Transient asystole was followed by consecutive ventricular escape beats, then transient bradycardia and approximately 5 seconds of complete heart block (atrio-ventricular, AV, dissociation, Fig. 7B2, RR and PP intervals plotted over time). As sinus rhythm returned, there was AV conduction (PR interval) variability in the first 7 beats, followed by stable atrial and ventricular rates and consistent PR intervals (Supplementary Fig. 6A-D). Exertional gasp movements of the chest were visible and detected on the ECG (Fig. 7B1). Video 1 illustrates the lethal event, which includes the pre-ictal period with epileptiform discharges, the onset of a GTCS, and the EEG/ECG manifestations during the post-ictal period and leading up to asystole/death. Following the lethal GTCS, Sudden Death Case 1 displayed post-ictal generalized EEG suppression (PGES, Fig. 7C2-5), apnea (based on video), inverted T-waves on the ECG (indicative of transient ischemia, Fig. 7C2), bradycardia (Fig. 7C2-5), atrial bigeminy (couplets of beats, Fig. 7C3), 2^nd^ degree AV block Type-2 (each QRS is preceded by a P-wave but QRS randomly drop, Fig. 7C4), 3^rd^ degree AV block (AV dissociation, Fig. 7C5) with focal ventricular activity, and terminal asystole. The rabbit went into rigor within 5 minutes following the lethal seizure.

**Fig. 7:**
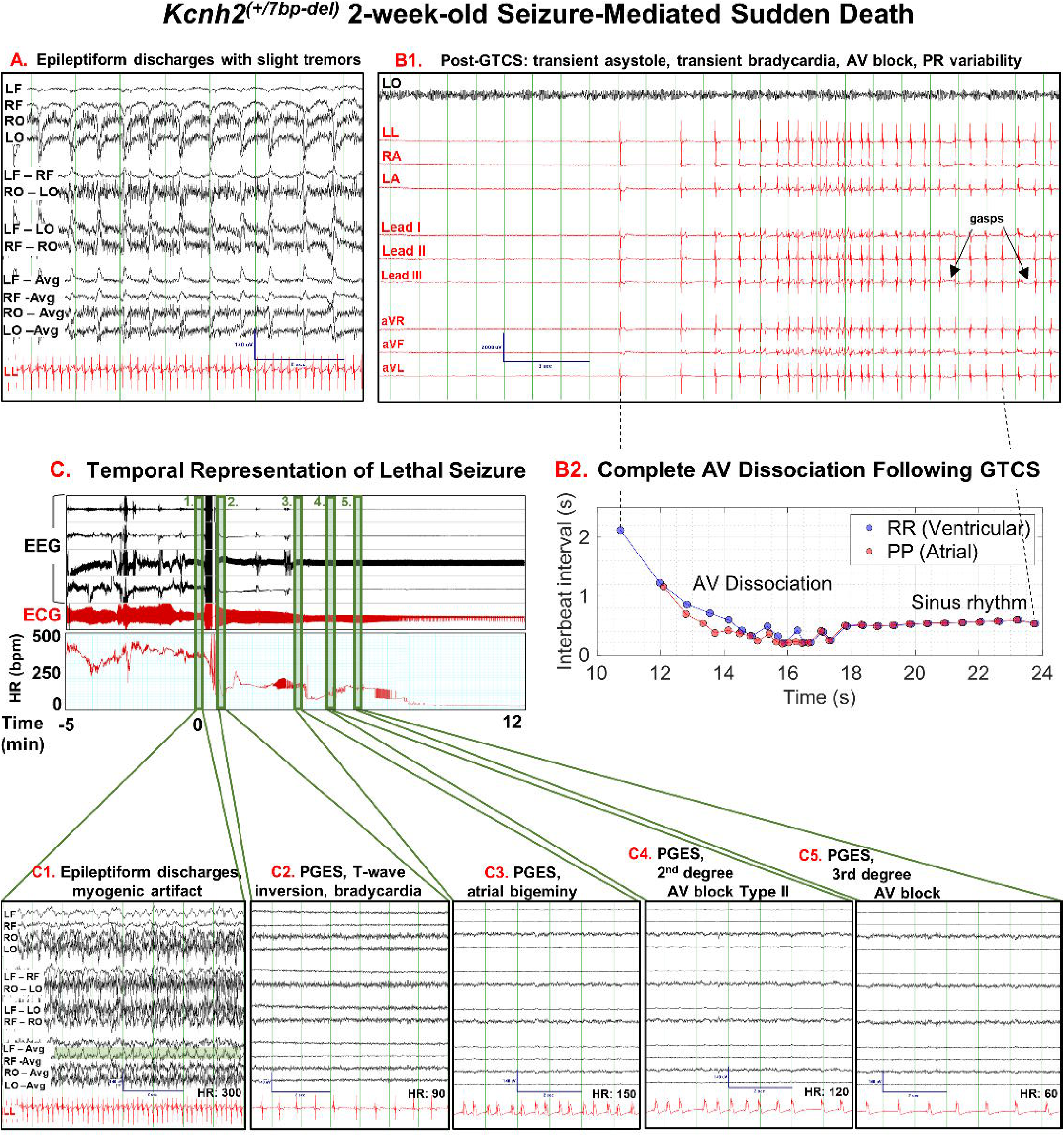
Sudden Death Case-1: Status-epilepticus-mediated sudden death in a 2-week-old female *Kcnh2*^(+/7bp-del)^ rabbit. **A.** Representative EEG (black) and ECG (red) traces showing epileptiform discharges and tremors ∼1.5 hours prior to sudden death. Scale bar shown for EEG signal: 140µV amplitude, 2 seconds. **B1.** Cardiac manifestations following a GTCS using one EEG trace, 3 referential ECG traces, and the standard bipolar and augmented ECG lead configurations (I, II, III, aVR, aVL, & aVF). Immediately following GTCS: sinus pause (∼8 seconds), transient bradycardia, AV block, and inverted T-waves. Rabbit was apneic for ∼19 seconds post-GTCS until first gasp (marked by arrow). Scale bar shown for ECG signal; 2000µV amplitude, 3 seconds. **B2.** Progression of cardiac abnormalities and recovery following a convulsive seizure. Ventricular rhythm with each QRS complex and atrial rhythm with each P wave plotted over time. **C.** Lethal seizure is depicted at minute zero. A cardiac tachogram is shown starting at 5 minutes pre-lethal seizure and ending close to asystole at 12 minutes post-lethal seizure. **1.** Pre-ictal interval shows epileptiform activity (highlighted in green), myogenic artifact from tremors, and heart rate at ∼300 BPM. Post-ictal intervals show **2.** PGES, bradycardia (HR: ∼90 BPM) and inverted T-waves, **3.** PGES and atrial bigeminy, **4.** PGES and 2^nd^ degree AV block, and **5.** PGES and 3^rd^ degree AV block. Scale bars shown for EEG signal; 140µV amplitude, 2 seconds.

Sudden Death Case 2 is a 1.8-month-old *Kcnh2*^(+/7bp-del)^ rabbit that suffered sudden cardiac death while collecting a routine 4mm ear biopsy for genotyping. Sudden stress is a known trigger for arrythmias in people with LQT2(21). Several cardiac pathologies were noted, which included atrial bigeminy, frequent periods of bradycardia (<96bpm), ECG ST- elevation/depression (indicative of ischemia, Supplementary Fig. 7A), premature ventricular complexes (PVCs), 2^nd^ degree AV block, and ultimately asystole. The rabbit also developed cardiogenic seizures. The clinical manifestations included myoclonic activity, however no epileptiform discharges were noted surrounding the convulsive seizures.

Sudden Death Case 3 is an 11-month-old female *Kcnh2*^(+/7bp-del)^ rabbit that died suddenly during a routine health check blood draw, likely also due to a stress-mediated response. The rabbit convulsed vigorously and went into rigor within 5 minutes of the lethal event (Table 1). Since there were not video-EEG-ECG recordings during this event, we cannot confirm whether the clinical manifestations were due to an epileptic or cardiogenic seizure.

Sudden Death Case 5 is a 21.7-month-old female *Kcnh2*^(+/7bp-del)^ rabbit that was found dead in the housing cage, 2 hours after last seen alive. The rabbit was found in full rigor and had a history of ECG abnormalities including a high incidence of PVCs (0.68/min, Supplementary Fig. 7B). No EEG abnormalities were noted in several recordings throughout the rabbit’s life.

Sudden Death Case 8 is a 2-month-old female WT rabbit that suffered a convulsive episode leading to death in the housing cage during the night. Continuous audio-video recording indicates episodes of tonic head extension and non-rhythmic dysynchronous forelimb motor manifestations prior to death, which is consistent with convulsive syncope. The rabbit was found deceased ∼7 hours post-mortem in full rigor. During a video/EEG/ECG recording ∼5 hours prior to sudden death, the rabbit exhibited PVCs (0.59/min), sinus pause (>2-sec), ST elevation, and deep inverted T-waves. In a recording taken 1 month prior to sudden death event, the rabbit exhibited several interictal epileptiform discharges.

In all other cases of spontaneous sudden death, the rabbits were found deceased unexpectedly in their housing cage and in full rigor (Table 1). In summary, there is a significantly higher prevalance of sudden death in *Kcnh2*^(+/7bp-del)^ vs. WT rabbits. While the cause of death was not definitive in all cases, none of the rabbits exhibited any findings during necropsy that would rule out SUDEP (e.g., structural abnormalities). In 3 cases of sudden death, we captured seizures and/or ECG abnormalities, which indicates the role of neuro-cardiac electrical abnormalities leading to spontaneous sudden death.

## Discussion

We developed the first translational knock-in model of *Kcnh2*-mediated LQT2. This study provides a comprehensive characterization of the molecular, biochemical, and *in vivo* neuro-cardiac phenotypes in *Kcnh2*^(+/7bp-del)^ rabbits, which mirror the cardiac and neuronal pathologies, and sudden death (i.e., SCD & seizure-mediated) seen in people with LQT2. There is a ∼50% reduction in total and WT *Kcnh2*, and full-length WT K_v_11.1 in *Kcnh2*^(+/7bp-del)^ rabbits. Consistent with loss-of-function *Kcnh2* variants(22, 23), there is a significant prolongation of ventricular repolarization in *Kcnh2*^(+/7bp-del)^ vs. WT rabbits. *Kcnh2*^(+/7bp-del)^ rabbits show an increased prevelance of spontaneous epileptiform discharges, electrographic seizures, and motor seizures, including myoclonic activity, clonic seizures, and GTCS. *Kcnh2*^(+/7bp-del)^ rabbits show an increased prevalence of spontaneous sudden death. These events include cases of SCD preceded by convulsive syncope, and seizure-mediated sudden death preceded by PGES and ECG abnormalities.

People with LQT2 are at an increased risk of both cardiac arrythmias and seizures(1, 9). Interestingly, 13% of SUDEP cases have variants associated with LQTS, including *KCNH2*(24). This translational model faciliates comprehensive assessments of the implications of *Kcnh2* variants in all organ systems, particularly the brain.

*Kcnh2*^(+/7bp-del)^ rabbits are more translationally relevant than cellular or rodent models, and faciliate studies that are not feasible in people. Heterologous cellular models expressing ion channel proteins of interest lack the native cellular machinery, accessory proteins, and physiological environment of the host(14). Expression is often at non-physiological levels, which generate results that are inconsistent with people(14). Human induced pluripotent stem cell-derived cardiomyocytes (hiPSC-CMs) are not fully mature, exhibit spontaneous activity, have a depolarized resting membrane potential, slow AP upstroke velocity, and the AP morphology more closely resembling fetal, rather than adult human cardiomyocytes(25). Cardiac ion channel expression, AP morphology, heart rate, and ECG parameters are very different in rodents vs. humans(14). Thus, the utility of mice as a model of cardiac arrhythmias and SCD remains controversial(26). While cardiac repolarization is driven by I_Kr_ in humans and rabbits, the transient outward and ultra-rapid potassium currents (I_to_ & I_Kur_) are the major repolarizing currents in rodents(27–29).

Several genetic mouse models of LQT2 have been generated with varying phenotypes(30–33). (1) Adult mice that overexpress the *Kcnh2-*G628S exhibit complete loss of I_Kr_. Yet, the mutant mice do not exhibit any changes in the AP duration, ECG intervals (e.g., QT_c_), or the susceptibility to pacing-induced arrhythmias(30). (2) Homozygous ERG1-B (*Kcnh2* isform B) knockout mice have no I_Kr_, but there is no difference in any ECG metrics in neonatal and adult mutant vs. WT mice(31).(3) Mice with homozygous deletion of the K_v_11.1 S4-S6 domain are embryonically lethal with abnormaliites in embryogenesis. Neonatal, but not adult, heterozygous mutant mice exhibit QT_c_ prolongation. (4) Mice homozygous for *Kcnh2*-N629D are embryonically lethal due to defects in cardiogenesis and vasculargenesis(32, 33). There is complete loss of I_Kr_ and prolonged AP durations in homozygous mutant myocytes. In contrast, 76.3% of myoyctes from heterozygous mutant embryos exhibit WT-like I_Kr_ and AP morphology(32). In contrast, juvenile and adult *Kcnh2*^(+/7bp-del)^ rabbits exhibit QT_c_ prolongation and SCD, and thus are a translational model of LQT2.

Several rodent models of epilepsy do not model the natural progression of seizure onset and are not physiologically relevant; these models require triggers to induce seizures(34, 35), have conditional, organ, or cell-type specific knock-out of genes of interest(36), do not reproduce the natural progression of clinical epilepsy, or have a high mortality rate that is not comparable to people with the type of epilepsy being modeled(37). Similar to our previous study that showed that 18% of people with LQT2 have a history of seizures/epilepsy, 19% of *Kcnh2*^(+/7bp-del)^ rabbits develop spontaneous epileptiform activity and seizures(9).

Rabbits provide a valuable model of cardiac arrhythmias and epilepsy, and are ideal for drug testing(29). Cardiac repolarization is driven by I_Kr_ in both humans and rabbits(29) . The cardiac electrical-activation recovery process is similar, as illustrated by similar AP and ECG morphologies, and sensitivity to I_Kr_ blockade(29). A cardiac-specific transgenic rabbit model of LQT2 exhibits prolonged QT and AP durations, reduced I_Kr_, polymorphic ventricular tachycardia, and high rate of SCD(38). However, these mutant rabbits have cardiac-specific overexpression of mutant human *Kcnh2*, in addition to the endogenous rabbit WT *Kcnh2*(38). This is in contrast to humans that only have 2 copies of each allele, *Kcnh2* expression is not driven by the beta-myosin heavy chain promoter and overexpressed in specific organs, and the genetic variant is present wherever the gene/protein is normally expressed. Importantly, the *Kcnh2*^(+/7bp-del)^ rabbits overcome limitations posed by cellular, rodent, or cardiac-specific transgenic rabbit models of LQT2. The CRISPR-Cas9-mediated knock-in 7bp frameshift deletion is in one allele of the endogenous rabbit *Kcnh2*, which better models the genetics seen in people with LQT2, and alters *Kcnh2*/K_v_11.1 expression and electrical function wherever *Kcnh2* is naturally expressed.

In the heart, loss-of-function variants in *KCNH2* cause cardiac AP and ECG-QT_c_ prolongation through a reduction in I_Kr_ density and/or alteration of the channel’s biophysical properties(39). The reduction in the repolarization reserve causes an imbalance between the cardiac depolarizing and repolarizing forces, leading to an increased risk of polymorphic ventricular tachycardia, ventricullar fibrillation, and sudden cardiac death(40). In the brain, pharmacological reduction in I_Kr_ depolarizes the resting membrane potential(41), reduces AP interspike intervals(42), and reduces spike-frequency adaption in neurons(42). Ultimately, this causes neuronal burst firing and hyperexcitability, which underlies epileptogenesis(43).

All rabbits that died suddenly were found in rigor (between 2-24 hours of last being seen alive). Rigor is due to ATP depletion, which can occur due to pre-mortem convulsive activity. We observed that in cases of pro-convulsant drug-induced seizure-mediated sudden death, there is a rapid onset of rigor in <30 minutes. This is in contrast to rabbits that are euthanized and do not go into rigor for >2 hours post-mortem. Rigor reaches 50% of maximum at >3.5-hours at 37°C, >6.5-hours at room temperature (17°C), and peak rigor is at >8-hours and >10-hours, respectively(44). Thus, many of the unwitnessed cases of sudden death were likley preceeded by vigorous muscle activity-mediated sudden depletion of ATP.

In a cohort of SUDEP cases that occured in epilepsy monitoring units, all deaths occurred following a generalized tonic-clonic seizure, which was accompanied by neuronal, cardiac, and respiratory dysfunction(45). Following the seizure, 90% of SUDEP cases demonstrated cardiorespiratory dysfunction, characterized by bradycardia, transient (>5 sec) or terminal apnea, and transient (>10 sec) or terminal asystole. Cardiac arrhythmias are a proposed mechanism of SUDEP(12). In patients with epilepsy, including those that later suffered SUDEP(46), peri-ictal arrythmias were recorded, including atrial fibrillation, asytole, supraventricular tachycardia, and bundle branch block(46, 47). Seizure duration was longer in patients with vs. without simultaneous cardiac abnormalities(47). Both long seizure duration, such as status epilepticus, and multi- system pathologies increase the risk of sudden death(45, 48). Ion channelopathies that increase the risk of both epilepsy and cardiac disease are found in SUDEP cases(24, 49). Variants in genes linked to Brugada syndrome (*SCN5A*), LQT3 (*SCN5A*), and Dravet syndrome (*SCN1A*) are reported in SUDEP cohorts(12). *KCNH2*-LOF-mediated LQT2, epilepsy, and sudden death was reported in a family with a heterozygous point mutation (c.246T>C, p.I82T)(50). The SUDEP case and twin sister showed generalized spike and slow wave EEG complexes, QT prolongation (550ms), and abnormal T-wave morphology. The *KCNH2* variant causes a significant decrease in I_Kr_ density and faster channel deactivation kinetics, confirming severe LOF(50).

This novel knock-in rabbit model of *Kcnh2*-mediated LQT2, epilepsy, and sudden death facilitates comprehensive studies to uncover the mechanisms underlying SCD and SUDEP in a population with both neuronal and cardiac electrical abnormalities.

## Limitations and Future Directions

Protein characterization was conducted using an antibody specific to full-length K_v_11.1^WT^. While this antibody demonstrates reduced K_v_11.1^WT^, which is in line with reduced WT *Kcnh2* expression, the antibody cannot detect mutant K_v_11.1 due to the frameshift and truncation. As many K_v_11.1 antibodies are raised in rabbits, particularly those upsteam of the 7bp-deletion, cross-reactivity prohibited assessments of total/mutant K_v_11.1 expression. While this manuscript did not examine the implications of the *Kcnh2* mutation on cellular electrophysiological properties, future studies will evaluate the changes in I_Kr_ and AP properties through comprehensive electrophysiological studies in HEK cells, cardiomyocytes, and neurons. This novel knock-in rabbit model of LQT2 will enable us to perform extensive molecular, cellular, and *in vivo* studies to better understand the implications of *Kcnh2* variants throughout the body. This model is a valuable tool to investigate the cardiac safety of medications in an at-risk disease population.

## Conclusions

We developed a novel translational genetic rabbit model of *Kcnh2*-mediated LQT2, which reproduces the neuro-cardiac electrical abnormalities and sudden death seen in people with LQT2. *Kcnh2*^(+/7bp-del)^ rabbits exhibit altered *Kcnh2*/Kv11.1 expression, cardiac QT_c_ prolongation, ECG abnormalities, epileptic seizures, and sudden death. Knock-in *Kcnh2*^(+/7bp-del)^ rabbits that demonstrate multi-system pathologies serve as a great tool to study the mechanisms for and cascade leading up to sudden death (e.g., SCD & SUDEP). *Kcnh2*^(+/7bp-del)^ rabbits provide a valuable translational model for the development of novel therapeutics, and cardiac safety drug testing.

## Supporting information

LQT2 Rabbit Lethal Event

## Declarations

*Ethics approval and consent to participate:* Not applicable

*Consent for publication:* Not applicable

*Availability of data and materials*: The data that support the findings of this study are available to qualified investigators through data transfer and user agreements upon request from the corresponding author.

*Competing interests*: The authors declare that they have no competing interests.

*Funding*: This work was supported by the National Institute of Health (DSA: NIH-NINDS 1R61NS133273); American Heart Association (DSA: 18CDA34110270); American Epilepsy Society (DSA: 060449-002); SUNY Upstate Pilot Grant; and the Department of Pharmacology at SUNY Upstate Medical University.

## Authors’ contributions

D.S.A contributed to the conception and design of the study. V.S, K.T.W, L.G.W, J.M, K.R.K, J.D.M, and R.S.G contributed to the acquisition and analysis of data. L.T.D and J.M.Z contributed to data validation. V.S, J.M.R, R.J.H.W, N.R.T, and D.S.A contributed to the preparation of the manuscript and figures.

## Acknowledgements

The work was supported by the National Institute of Health (DSA: NIH-NINDS 1R61NS133273), American Heart Association (DSA: 18CDA34110270), American Epilepsy Society (DSA: 060449-002), Upstate Medical University Pilot Grant, and the Department of Pharmacology at SUNY Upstate Medical University. Dr. Joseph Miano was instrumental in designing the molecular tools for the CRISPR-Cas9 genetic reprograming.

## List of Abbreviations

LQTS: Long QT Syndrome
LQT1: Long QT Syndrome Type-1
SCD: Sudden Cardiac Death
SUDEP: Sudden Unexpected Death in Epilepsy
ECG: Electrocardiogram
ONT: Oxford Nanopore Technology
LQT2: Long QT Syndrome Type-2
GTCS: Generalized tonic-clonic seizures
LOF: Loss-of-function
AV: Atrio-ventricular
AP: Action Potential
PGES: Postictal Generalized EEG Suppression
EEG: Electroencephalogram
PVC: Premature Ventricular Complex

