## Supplemental Methods for "Knock-in *Kcnh2* Rabbit Model of Long QT Syndrome Type-2, Epilepsy, and Sudden Death"

**Supplementary Methods**

*Tissue Collection:* Following sudden death or euthanasia (Euthaphen, Dechra Veterinary Products) a necropsy is performed on all rabbits, which includes a meticulous dissection and collection of eight brain regions (frontal cortex, parietal cortex, temporal cortex, occipital cortex, cerebellum, hippocampus, pons, and medulla), and seven heart regions (right atrium, right ventricle, left atrium, left ventricle, left ventricular endocardium, ventricular apex, and interventricular septum). All tissue is collected in labeled tubes, flash frozen in liquid nitrogen, and stored at -80ºC.

*Genotyping* (All primer and probe sequences are listed in Supplementary Table 1): Genomic DNA (gDNA) is isolated from rabbit ear tissue (10-15 mg) using the Qiagen DNeasy Blood and Tissue Kit and amplified through PCR using a *Kcnh2*-specific forward primer (amplify total *Kcnh2*, loading control) and a 7bp-del mutation-specific forward primer. The mutation-specific primer flanks the site of the deletion and binds to the *Kcnh2* 7bp-del variant allele. A common reverse primer is used to amplify both total and mutant *Kcnh2*, which is 109bp downstream of the deletion site. PCR products are visualized by electrophoresis on a 3% agarose gel with 1x Tris-acetate EDTA (TAE) running buffer at 150 V for 10 min, then at 80 mA for 60 min. Sanger sequencing is performed to confirm the genotype.

*RNA Sequencing of the Rabbit Genome:* We examined total and allele-specific expression of *Kcnh2* using a combinatorial approach of genome-wide RNA sequencing, targeted long-read RNA sequencing, and qPCR. To build the rabbit genome, the OryCun 2.0 genome assembly and OryCun 2.0.112 annotations obtained from Ensembl are used as input to the genomeGenerate function in STAR (v2.7.11b); sjdbOverhang is set to 149. Single reads were aligned to the genome. The featureCounts function as part of the Subread package (v2.0.6) was used to obtain gene counts from BAM files. Gene counts are scaled to 1 million reads for each sample. Rsamtools (v2.20.0) and GenomicAlignment (v1.40.0) packages are used to query bam files to quantify gapped reads. Total reads showing at least 70% matching among the 5 bases on either end of the 7bp deletion are counted as spanning the site of the deletion, of which, those showing gapped alignments at the site of the 7bp deletion are counted as gapped reads. The ratio of gapped to total spanning reads quantifies the proportion of transcripts that are positive for the 7bp deletion.

*Oxford Nanopore Technology (ONT) Sequencing & Quantitative PCR (qPCR):* RNA is extracted from rabbit tissue (45-50 mg) using the Zymo DNA/RNA Shield and Zymo Quick RNA Miniprep Kit. cDNA is generated using the BioRad iScript cDNA Synthesis Kit. For ONT sequencing, specific primers are used to amplify a region that surrounds the region of the mutation, which yields a 636bp product in WT *Kcnh2* cDNA and a 629bp product in the 7bp-del *Kcnh2* cDNA; the primers yield a 1100bp product in WT *Kcnh2* gDNA and a 1093bp product in the 7bp-del *Kcnh2* gDNA (see Online Supplement Table 1 for sequences/probes). Plasmidsaurus provides a read-by-read assessment of WT vs. mutant alleles, which are counted in order to determine the ratio of each allele in the mutant tissue. Importantly, in contrast to qPCR and RNAseq, this does not provide an estimate of total *Kcnh2* abundance.

For qPCR, each primer binds to a separate exon to ensure that only cDNA is amplified. The primers yield a 216bp product in WT *Kcnh2* and a 209bp product in 7bp-del *Kcnh2*. Sanger sequencing is performed for all primer pairs to confirm the amplification of both alleles in the region of interest, and that the probes are specific for WT vs. mutant sequences. Standard curves are generated using known concetrations of pure synthetic DNA (gblocks) to determine the relationship between the Ct value and the copy number of the input sample. qPCR analyses are performed in triplicates on each of the 15 brain/heart regions from 14 animals, in parallel with a 6-point standard cDNA concentration curve run in triplicate. Each experimental sample is tested seperately with a WT-specific *Kcnh2* probe, mutant-specific *Kcnh2* probe, and *Actb* amplification. The copy number of each *Kcnh2* sample is normalized to the copy number of *Actb* of the same sample. As we ran a total of 630 samples, alongside the six-point standard curves, we utilized two 384-well plates. To adjust for inter-plate variablity, all experimental samples are normalized to the average of the WT *Kcnh2:Actb* ratio from all WT apex samples on that plate (Eq. 1).

$\frac{Experimental Sample Ct}{\frac{WT Apex Kcnh2 Ct}{WT Apex Actb Ct}}$	(Eq. 1)

*Novel Generation of Anti-K_v_11.1 Antibody:* Antisera are raised in two Guinea pigs (GP5 and GP6) against a peptide (Supplementary Table 1) corresponding to the C-terminus of rabbit K_v_11.1, which is completely conserved in mice, rats and humans. The peptide is synthesized with an additional C at the N-terminus to allow for coupling to keyhole limpet hemocyanin. Coupling, injections, animal care and exsanguination are performed by Cocalico Biological, Inc (Stevens, PA). GP6 serum is the most immunoreactive and is affinity purified using the same peptide coupled to SulfoLink columns as previously described^1^.

*Exogenous Kcnh2 Expression:* Plasmid constructs in pCDNA3.1(+) are generated using the published rabbit *Kcnh2* sequence (NM_001082384.1) for the WT contruct and a 7bp deletion (1627bp-1633bp, NM_001082384.1) for the 7bp mutant construct (Invitrogen Geneart Gene Synthesis, ThermoFisher). For assessments of protein expression, we generated MYC-WT-K_v_11.1 and FLAG-7bp-del-K_v_11.1 construsts; the tags are at the N-terminus of K_v_11.1. HEK293T cells are maintained at 37ºC and 5% CO_2_ in high glucose Dulbecco’s modified Eagle’s growth medium supplemented with 10% fetal bovine serum, 100µg/mL penicillin, and 100µg/mL streptomycin. HEK293T cells are seeded into 6-well plates at a concentration of 6 x 10^5^ cells/well. Cells are transfected 24 hours post-seeding using 6µL of 1 mg/mL PEI and 0.5 µg of cDNA of interest. Empty vector pcDNA3.1(+) is used as the negative control.

*Cell lysis, Tissue Homogenization, SDS-Page, and Western Blot:* Cells are harvested 20 hours post-transfection using HBSE and lysed in ice-cold Triton lysis buffer (150 mM NaCl, 50 mM Tris-HCl, 1 mM EDTA, 1% Triton X-100, 10 μM pepstatin A, 0.2 μM soybean trypsin inhibitor, 0.2 mM phenylmethylsulfonyl fluoride, 1 mM DTT, pH 8.0) on ice for 30 min. Lysates are spun down at 16,000xg for 10 mins at 4ºC. All sample pellets are resuspended in gel-loading buffer^2^ incubated at 37°C for 30 min, and subjected to SDS-PAGE by loading 20 µg protein for each sample.

The tissue is weighed to achieve equal amounts of tissue from each brain/heart region, which is used for immunoblotting. Brain tissue samples are prepared using a dounce homogenizer in homogenization buffer (10mM Tris-Base, 1mM EGTA, 10 μM pepstatin A, 0.2 μM soybean trypsin inhibitor, 0.2 mM phenylmethylsulfonyl fluoride, 1 mM DTT, pH 7.4). Heart tissue samples are prepared with homogenization buffer using a polytron. Following homogenization, samples are spun at 300xg for 10 minutes at 4ºC. The supernatant is removed and spun again at 18,000xg for 10 minutes at 4ºC. The pellets are collected and resuspended in additional homogenization buffer and the protein concentration is measured with the Bradford method. All samples are resuspended in gel-loading buffer, boiled at 98ºC for 3 minutes, then subjected to SDS-PAGE by loading 15µg protein per sample. Proteins are transferred to nitrocellulose^3^ and probed with our in-house K_v_11.1, MYC (#2276, Cell Signaling Technology), FLAG (clone M2 #F3165, Sigma), Na/K ATPase α1 (#3010, Cell Signaling Technology), and GAPDH (anti-GAPDH #G2320, Santa Cruz) antibodies in TBS blocking buffer (0.1% Tween, 4% BSA, 0.1% Sodium Azide), Na/K ATPase and GAPHD serve as loading controls. Immunoreactivity is detected using Pico Chemiluminescent Substrates (ThermoFisher #34579) and a Bio-Rad ChemiDoc imager.

*In-Vivo Video-ECG-EEG Signal:* Recordings are acquired from male and female rabbits throughout their lifespan, which included pre-pubescent (<3-months of age) and fertile (>6-months of age) rabbits. A total of 9 subdermal clinical grade bent (chest) and 13mm straight (scalp) pin electrodes (Rythmlink, Columbia, South Carolina) are used to acquire acute video-ECG-EEG recordings. Head (EEG): 4 pin electrodes are placed at the left frontal, right frontal, left occipital, and right occipital regions, with a 5^th^ pin placed in the center at clinical EEG position Cz that serves as the reference. Chest (ECG): 3 pin electrodes are placed in the axilla near the left arm, right arm, and left leg, with the ground placed in the axilla near the right leg. All data is acquired using the NATUS Quantum amplifier and Neuroworks v9.3.1 software at a sampling rate of 1024 Hz.

*ECG Analysis:* Analysis includes the generation of 40-beat averages, which are manually adjudicated throughout the entire 5-min epoch, to calculate the heart rate (HR), cardiac conduction (P, PR, & QRS), and cardiac repolarization metrics (QT, JT_end_, and T_peak_-T_end_). JT_peak_ is manually calculated by substracting T_peak_-T_end_ from JT_end_. QT, JT_end_, and JT_peak_ values are corrected for heart rate using a rabbit-specific formula^4^ (Eq. 2). JT_peak_ and JT_end_ were corrected using the same formula (JT_pc_ & JT_ec_). Figure 3C illustrates each of the ECG metrics.

$\mathrm{QT}_{c}=\frac{\mathrm{QT}}{10 \times\mathrm{RR}^{0.72}}$	(Eq. 2)

*Statistical Analysis:* Continuous data is tested for normality using Komogorov-Smirnov test. If normally distributed, t-test (for two groups) or ANOVA (for more than two groups) is used. If the data is not normally distributed, Mann-Whitney/Wilcoxon RankSum test (two groups), or Kruskal-Wallis (>2 groups) is used. Logistic regression is used to adjust for multiple variables (age, sex, & heart rate) when testing for differences in ECG metrics. Chi-squared or Fisher’s exact test is used to test for differences between proportions.
