## Supplemental Figures for "Knock-in *Kcnh2* Rabbit Model of Long QT Syndrome Type-2, Epilepsy, and Sudden Death"

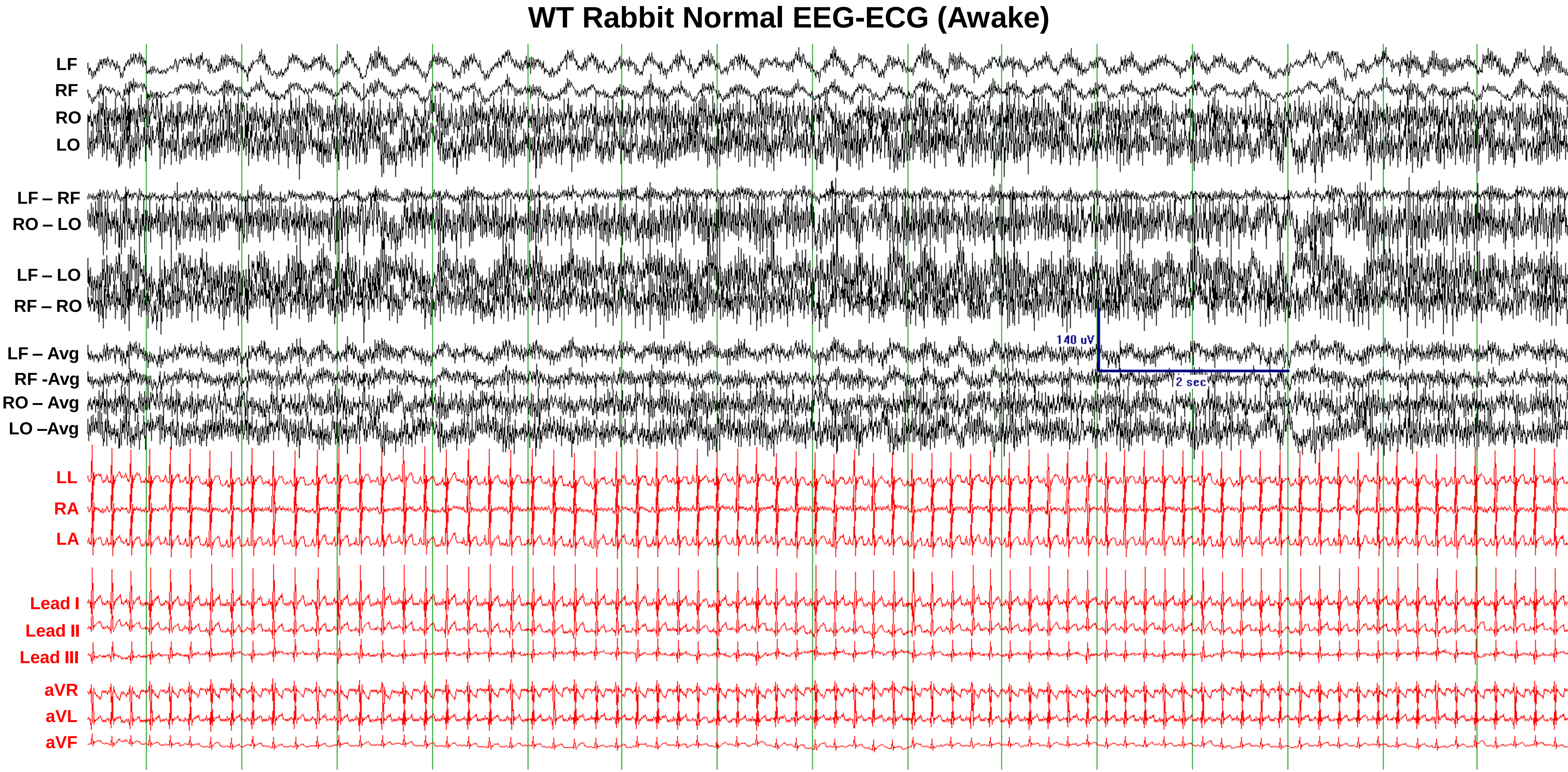


**Supplementary Fig. 1:** Baseline EEG (black) and ECG (red) in a 1.5-month-old male WT rabbit. EEG: Referential, bipolar, and unipolar traces. ECG: Referential, bipolar limb (I, II, III), and augmented (aVR, aVL, aVF) lead configurations. Scale bar indicates EEG=140μV and 2 seconds.

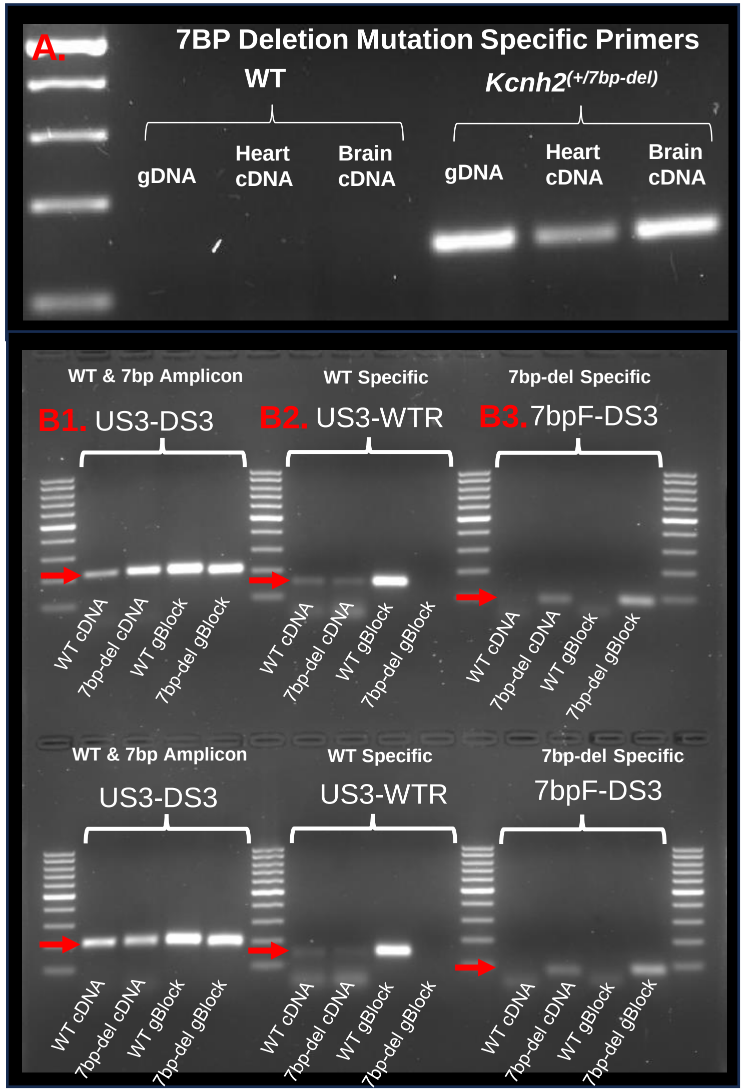


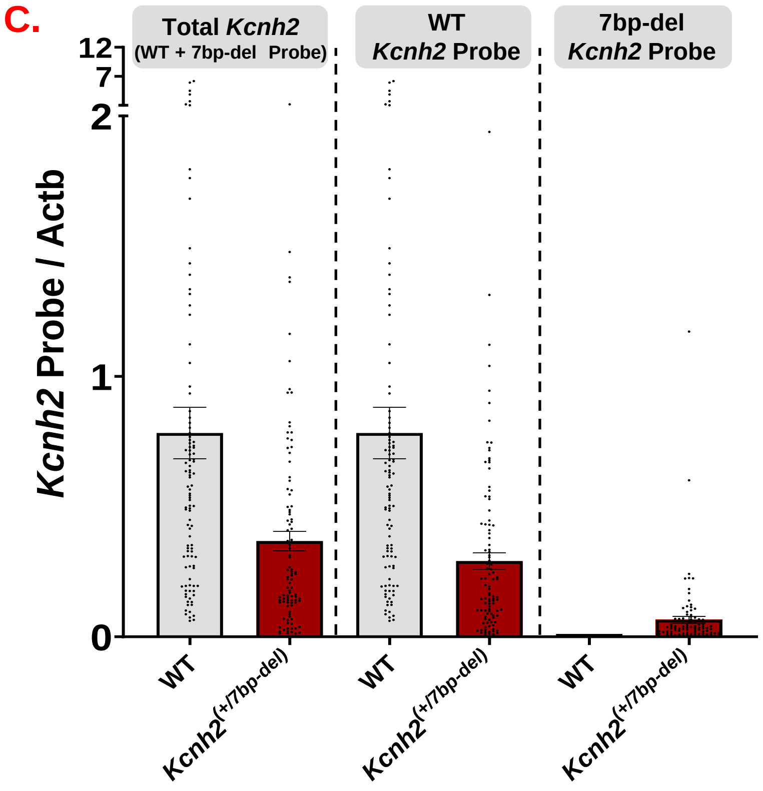


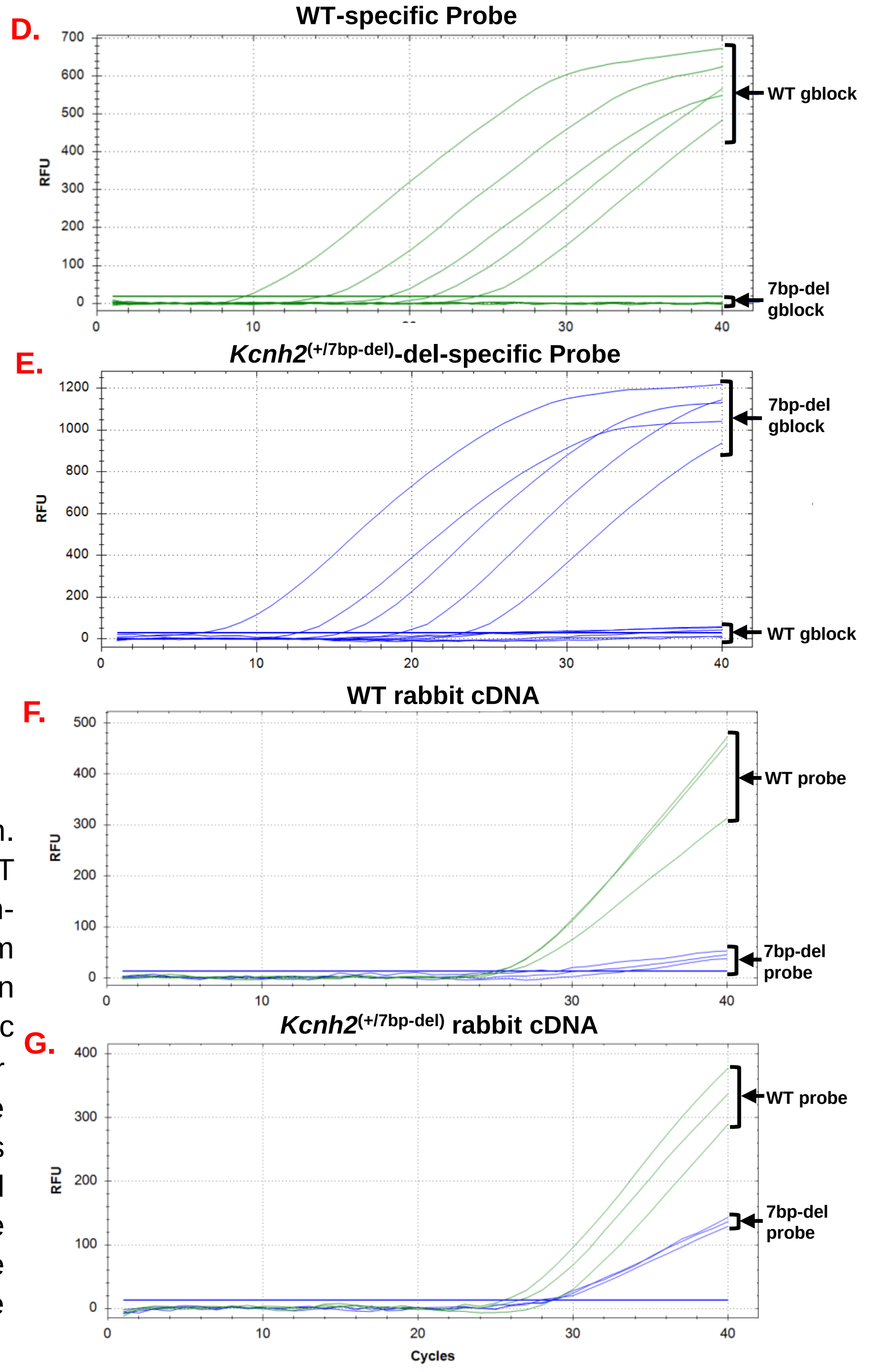


**Supplementary Fig. 2:** qPCR primer and probe verification. **A.** PCR of genomic DNA and complimentary DNA of the WT and *Kcnh2*$^{(+/7bp\text{-}del)}$ heart and brain tissue, using a mutation-specific primer. **B1.** *Kcnh2* upstream (US3) and downstream (DS3) primers amplify 216bp region in WT and 209bp region in 7bp-del cDNA and synthetic DNA (g-block). **B2.** WT-specific primer amplifies 162bp region in WT and 7bp-del cDNA and WT synthetic DNA. **B3.** 7bp-del-specific primer amplifies 89bp region in 7bp-del cDNA and synthetic DNA. WTR: WT-reverse; 7bpF: 7bp-del-forward. **C.** Relative gene expression of total *Kcnh2* (WT + mutant), WT *Kcnh2*, and 7bp-del mutant *Kcnh2* transcripts grouped across fifteen regions in WT and mutant tissue (WT: N=7 rabbits; 7bp: N=7 rabbits). **D.** Amplification plots of probe-based qPCR. WT-specific probe with WT synthetic DNA and 7bp-del mutant synthetic DNA. **E.** 7bp-del-specific probe with WT synthetic DNA and 7bp-del mutant synthetic DNA. **F.** cDNA from WT rabbit tissue with WT-specific probe (green) and 7bp-del-specific probe (blue). **G.** cDNA from *Kcnh2*$^{(+/7bp\text{-}del)}$ rabbit tissue with WT-specific probe (green) and 7bp-del-specific probe (blue).

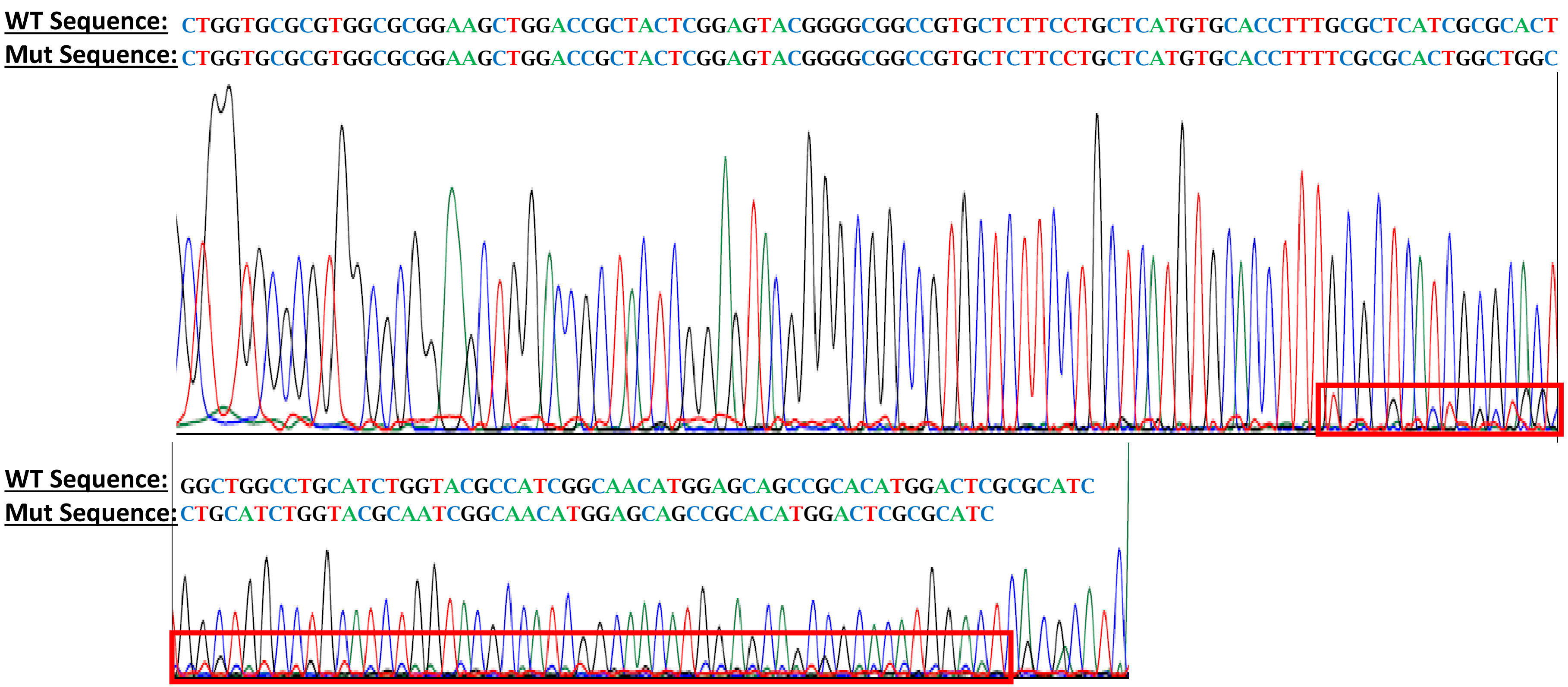


**Supplementary Fig. 3:** Sanger Sequencing Electropherogram. qPCR primers used in *Kcnh2*$^{(+/7bp\text{-}del)}$ cDNA. Primers used: US3, DS3.

| **Primer Description (NM_001082384.1)** | **Primer Sequence** | **Location** |
|---|---|---|
| ***Genotyping Primers*** | | |
| **Rabbit *Kcnh2*-specific forward primer (GenF1)** | 5’ ctggcgtgggacgtgcttca 3’ | Intron 5 |
| **Total and mutant rabbit *Kcnh2* common reverse primer (GenR1)** | 5’ aggccgctgctgttgtagg 3’ | Exon 7 |
| **7bp-specific rabbit *Kcnh2* forward primer** | 5’ ttgctcatgtgcaccttttcgcg 3’ | Exon 7 |
| ***qPCR Primers*** | | |
| **Forward primer for rabbit *Kcnh2* (US3)** | 5’ atcttcggctctggctctgag 3’ | Exon 6 |
| **Reverse primer for rabbit *Kcnh2* (DS3)** | 5’ gatgcgcgagtccatgtg 3’ | Exon 7 |
| **Forward primer for rabbit *Actb*** | 5’ atgtgcaaggccggctt 3’ | Exon 2 |
| **Reverse primer for rabbit Actb** | 5’ acgatgccgtgctcgat 3’ | Exon 3 |
| ***ONT Sequencing Primers*** | | |
| **Forward primer for rabbit *Kcnh2*** | 5’ ctgccttcttgctgaag 3’ | Exon 6 |
| **Reverse primer for rabbit *Kcnh2*** | 5’ agaagatcttctccgagttg 3’ | Exon 7 |
| **Probe Description** | **Probe Sequence** | |
| ***ONT Sequencing Probes*** | | |
| **WT-specific Kcnh2 probe sequence** | /56-FAM/agccagtgc/ZEN/gcgatgagc | |
| **Mutant-specific (7bp) Kcnh2 probe sequence** | /5HEX/ ttgctcatg/ZEN/tgcaccttttcgcg | |
| ***$K_v$11.1 Antibody*** | | |
| **$K_v$11.1 C-terminus epitope amino acid sequence** | CGALTSQPLHRHGSDPGS | |

**Supplementary Table 1:** Sequences of all primers and probes used for PCR, qPCR, and ONT sequencing. Amino acid sequence of anti-$K_v$11.1.

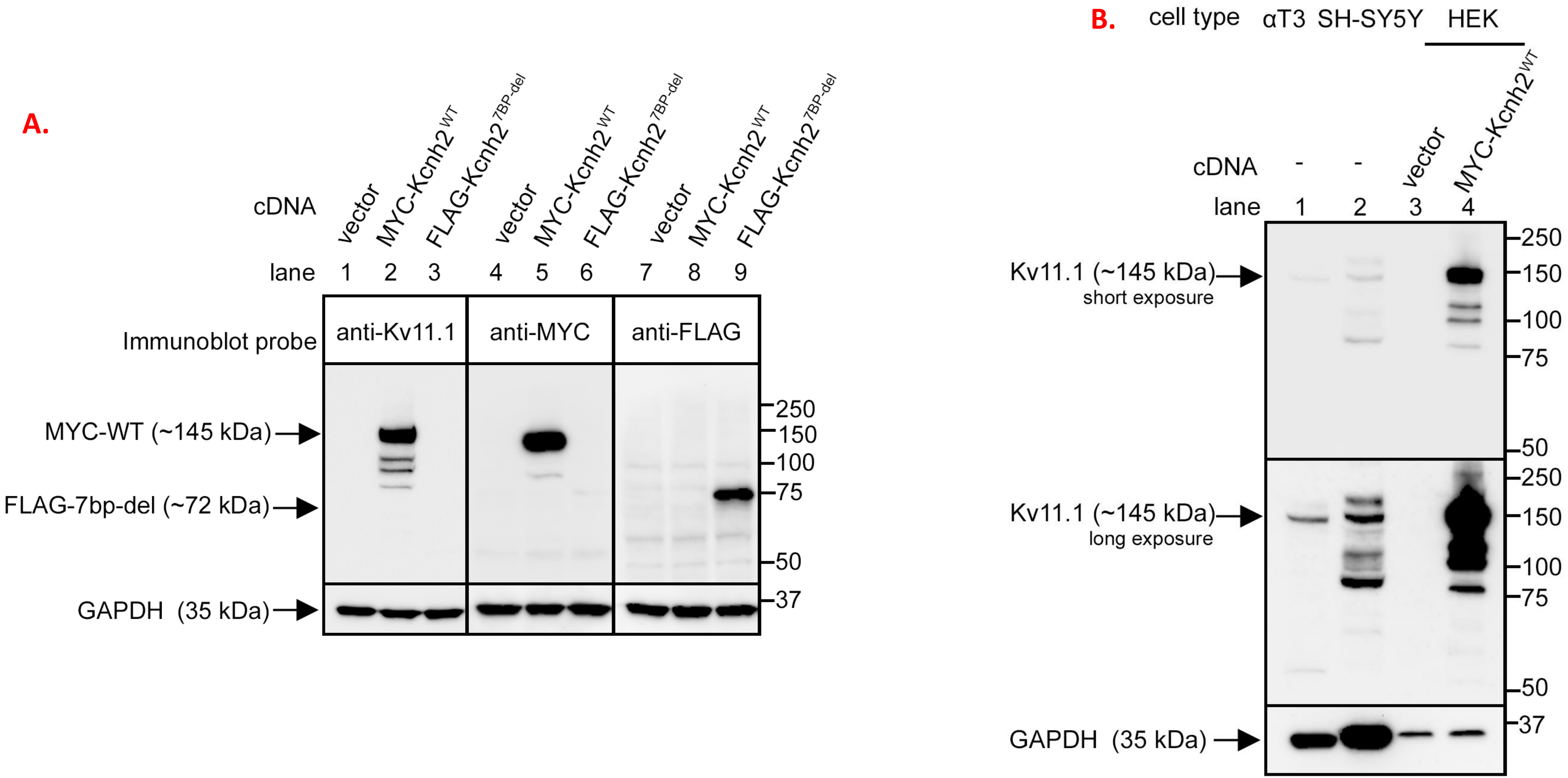


**Supplementary Fig. 4: A.** Immunoblot showing immunoreactivity of anti-$K_v$11.1 and anti-MYC in recognizing myc-$K_v11.1^{WT}$ and anti-FLAG in recognizing $K_v11.1^{7bp\text{-}del}$ constructs over-expressed in HEK cells. **B.** Immunoblot showing immunoreactivity of anti-$K_v$11.1 in recognizing endogenously expressed $K_v$11.1 in mouse pituitary (α-T3) and human neuroblastoma (SH-SY5Y) cell lines.

| | Wild Type | 7bp-del | Statistical Analyses | | |
|---|---|---|---|---|---|
| | | | Wilcoxon Rank Sum | Logistic Regression | |
| **ECG Measure** | **Mean ± Standard Deviation** | **Mean ± Standard Deviation** | **p-value** | **Odds Ratio (95% CI)** | **p-value** |
| **Heart Rate** | 273.91 ± 52.052 | 287.04 ± 53.10 | 0.164767396 | 0.988 (0.979,0.997) | **0.010008996** |
| **P Duration** | 25.47 ± 4.37 | 26.08 ± 5.48 | 0.891218847 | 0.967 (0.863,1.082) | 0.554669121 |
| **PR** | 59.21 ± 10.23 | 54.84 ± 8.56 | **0.018110803** | 1.099 (1.019,1.185) | **0.014812847** |
| **$QT_c$** | 244.51 ± 18.82 | 279.14 ± 21.75 | **$2.77743 \times 10^{-14}$** | 0.913 (0.882,0.944) | **$1.5225 \times 10^{-7}$** |
| **QRS** | 25.09 ± 5.21 | 24.84 ± 5.09 | 0.605954274 | 1.022 (0.899,1.161) | 0.742333248 |
| **$JT_{ec}$** | 193.98 ± 16.64 | 227.41 ± 19.03 | **$1.95451 \times 10^{-15}$** | 0.895 (0.86,0.932) | **$5.79916 \times 10^{-8}$** |
| **$JT_{pc}$** | 137.91 ± 21.91 | 180.46 ± 19.51 | **$3.22658 \times 10^{-15}$** | 0.919 (0.893,0.947) | **$1.88463 \times 10^{-8}$** |
| **$T_pT_e$** | 27.74 ± 9.57 | 22.40 ± 5.86 | **0.002081525** | 1.104 (1.037,1.174) | **0.00188449** |

**Supplementary Table 2:** Conduction and repolarization ECG metrics in WT and *Kcnh2*$^{(+/7bp\text{-}del)}$ rabbits. Wilcoxon rank sum test; logistic regression model adjusts for age, sex, and heart rate.

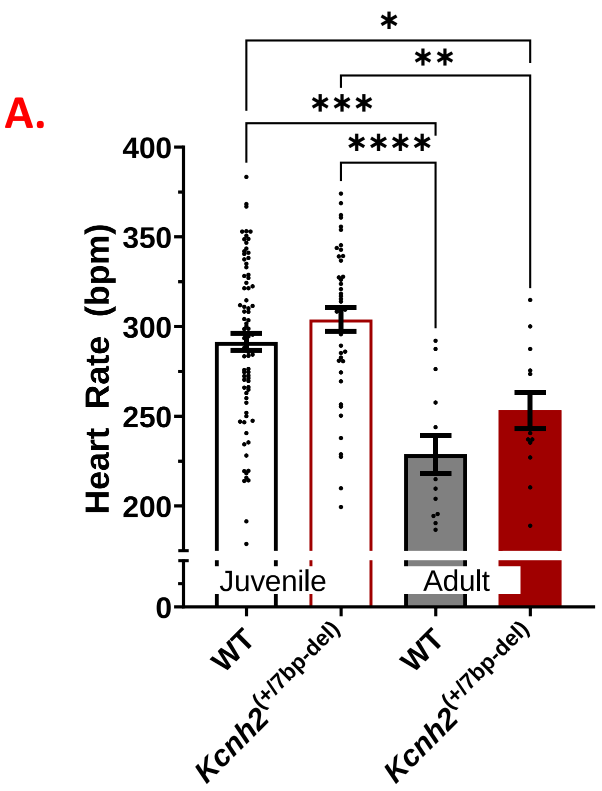


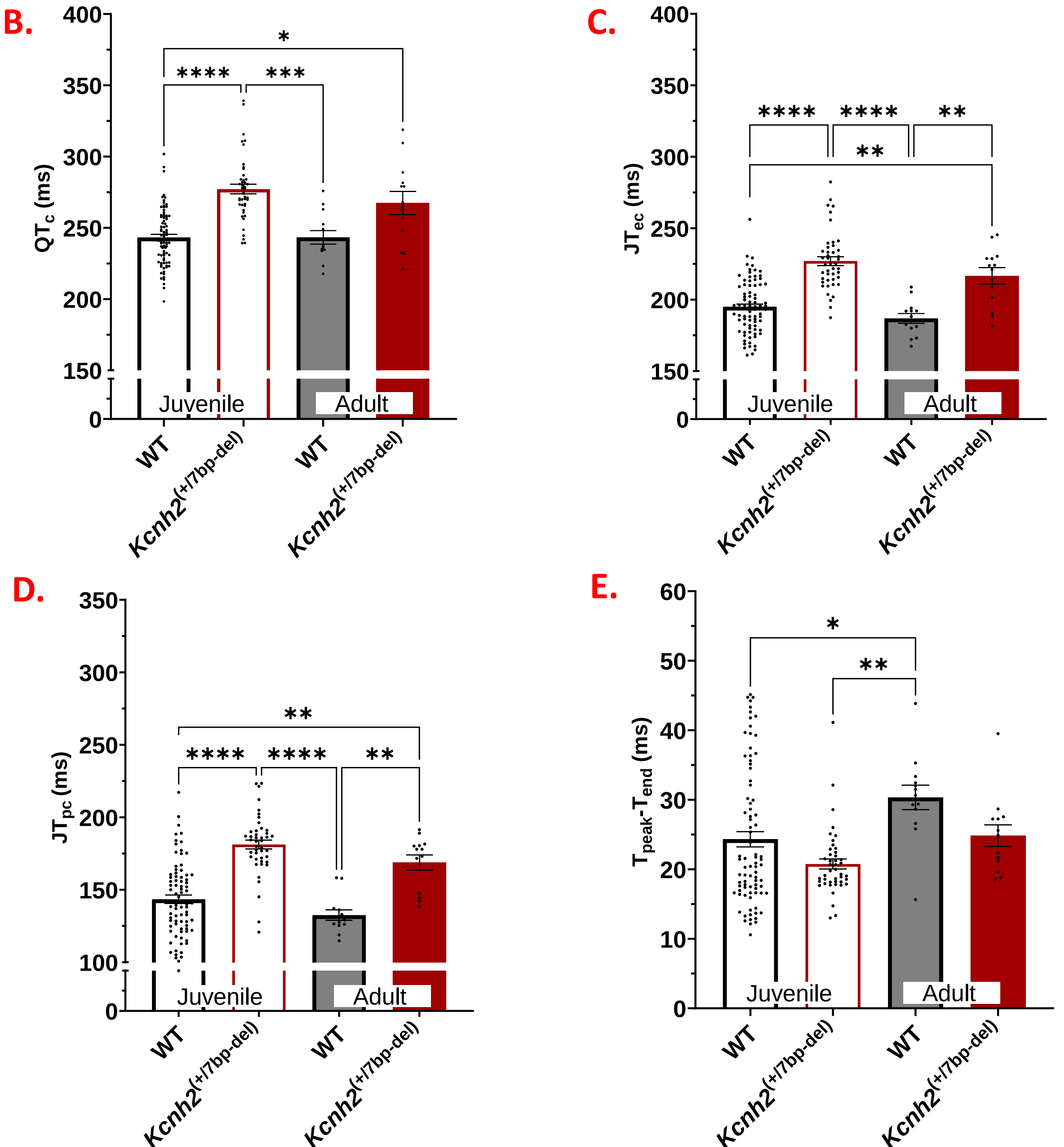


**Supplementary Fig. 5:** Genotype-specific ECG measures at juvenile (<3 months) and adult (>6months) timepoints. Each point is a 5-minute period of the recording. WT (N=68, n=89), *Kcnh2*$^{(+/7bp\text{-}del)}$ (N=37, n=52).

| A. $QT_c$: Baseline (All Littermates) | | | | |
|---|---|---|---|---|
| | **N** | **Median (ms)** | **Standard Deviation (ms)** | **Wilcoxon p-value** |
| **WT** | 2 | 263.3 | 21.6 | <0.0001 |
| **7bp-del** | 4 | 310.7 | 7.4 | |

| B. $QT_c$: Percent change in $QT_c$ in the *Kcnh2*$^{(+/7bp\text{-}del)}$ Sudden Death Case-1 on the lethal vs. non-lethal days | | | |
|---|---|---|---|
| | **Median (%)** | **Standard Deviation (%)** | **Signed-Rank p-value** |
| **7bp-del** | 10.3 | 7.2 | <0.0001 |

**Supplementary Table 3:** Cardiac $QT_c$ duration in Sudden Death Case-1 and its littermates: **A.** Baseline $QT_c$ recorded 2-days prior to sudden death in 3 *Kcnh2*$^{(7bp\text{-}del)}$ (one is Sudden Death Case 1) and 2 WT littermates. **B**. Percent change in $QT_c$ in the *Kcnh2*$^{(+/7bp\text{-}del)}$ Sudden Death Case-1 on the lethal vs. non-lethal days.

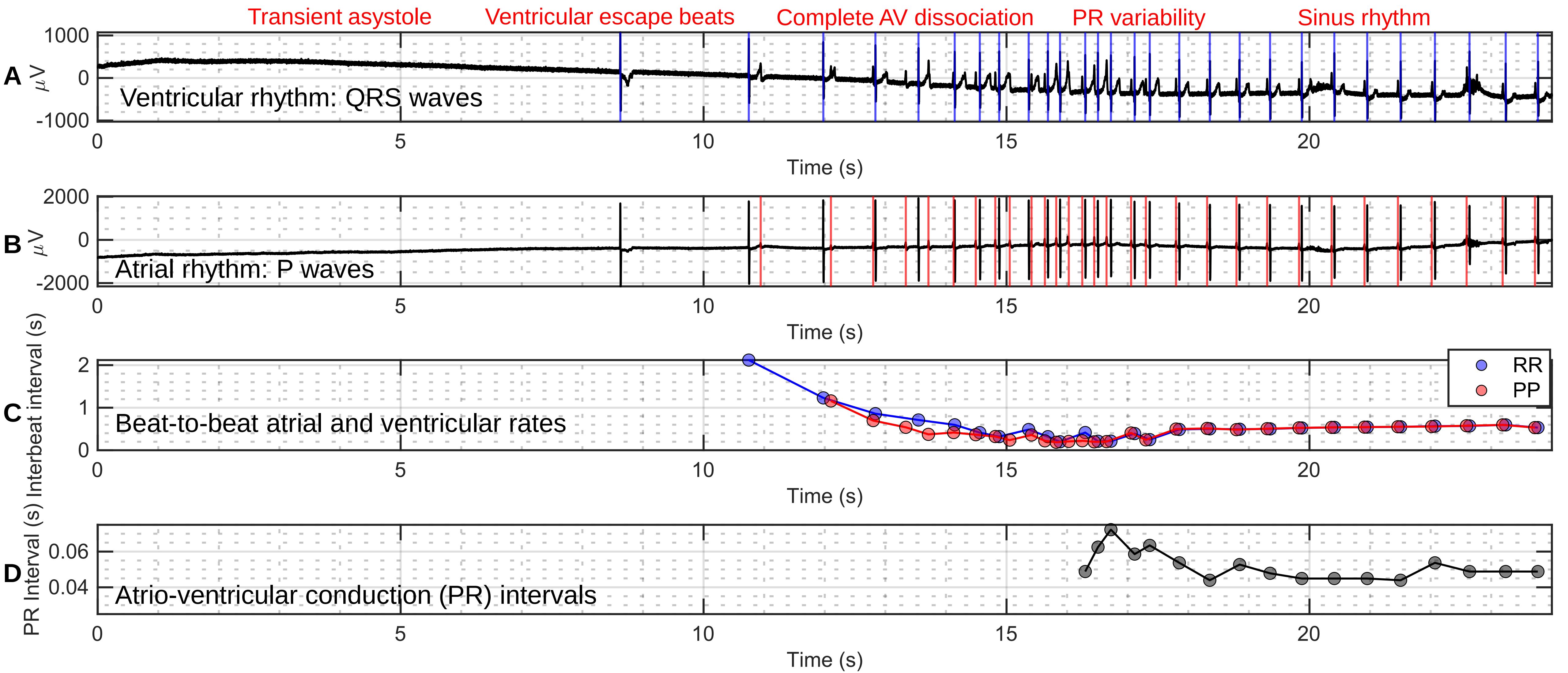


**Supplementary Fig. 6:** Progression of cardiac abnormalities and recovery following a convulsive seizure. **A.** Ventricular rhythm: ECG referential lead LL with each QRS complex annotated. **B.** Atrial rhythm: ECG referential lead LA with each P wave annotated. **C.** Instantaneous rates: Ventricular (RR) and atrial (PP) beat-to-beat intervals. **D.** Atrio-ventricular conduction: PR intervals for all sinus beats plotted over time. LL: left leg, LA: left arm.

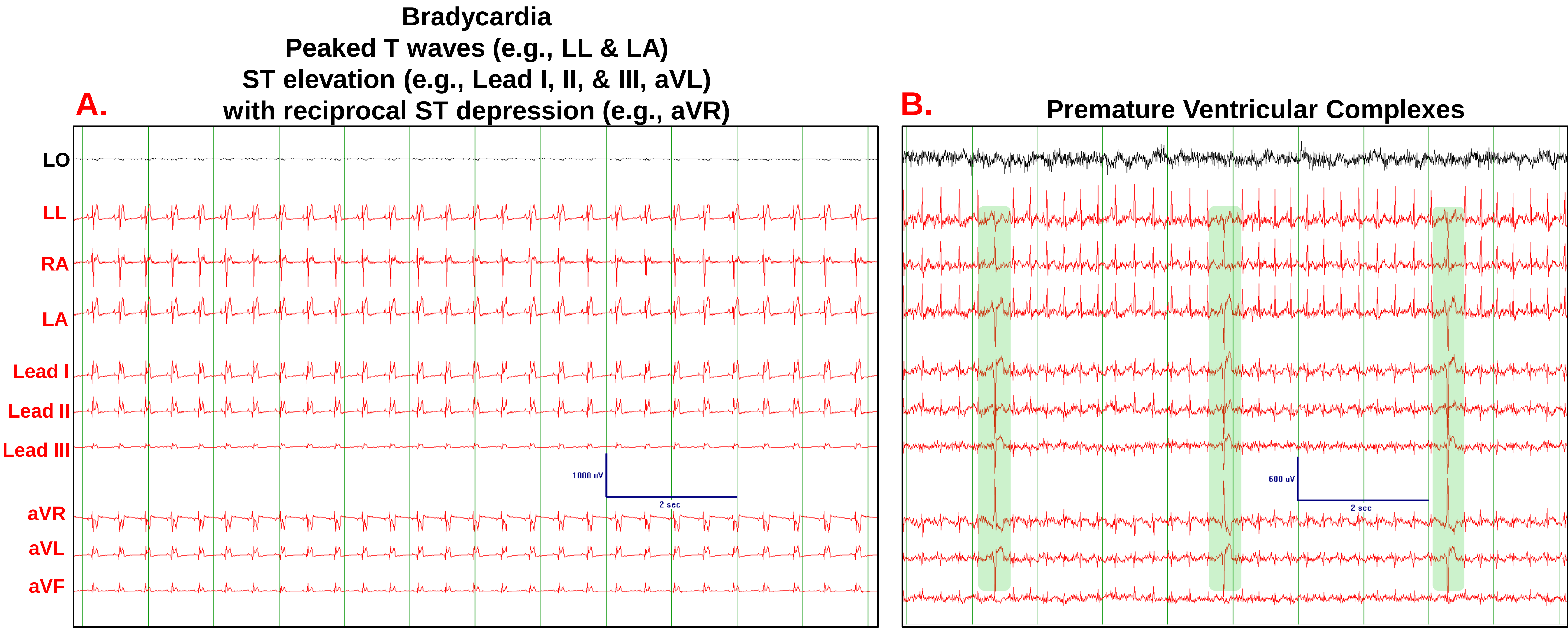


**Supplementary Fig. 7:** Cardiac abnormalities noted in *Kcnh2*$^{(+/7bp\text{-}del)}$ rabbits, depicted using referential and standard (I, II, III, and augmented) ECG lead configurations. **A.** Peaked T waves in referential leads (LA, LL). ST elevation in leads I, II, and aVL and reciprocal ST depression in aVR. Sudden death Case 2: *Kcnh2*$^{(+/7bp\text{-}del)}$ male 7-week-old rabbit **B.** PVC: Premature Ventricular Complex. Sudden Death Case 5: *Kcnh2*$^{(+/7bp\text{-}del)}$ female 13-month-old rabbit. ECG scale bar indicates 1000μV (A) and 600μV amplitude (B) and 2 seconds.
